# The propofol binding sites of prokaryotic voltage-gated sodium channels

**DOI:** 10.1101/2020.10.28.359513

**Authors:** Elaine Yang, Weiming Bu, Antonio Suma, Vincenzo Carnevale, Kimberly C. Grasty, Patrick Loll, Kellie Woll, Natarajan Bhanu, Benjamin A. Garcia, Roderic G. Eckenhoff, Manuel Covarrubias

## Abstract

Propofol, one of the most commonly used intravenous general anesthetics, modulates neuronal function by interacting with ion channels. The mechanisms that link propofol binding to the modulation of distinct ion channel states, however, are not understood. To tackle this problem, we investigated prokaryotic ancestors of eukaryotic voltage-gated Na^+^ channels (Navs) using unbiased photoaffinity labeling with a photoacitivatable propofol analog (AziP*m*), electrophysiological methods and mutagenesis. The results directly demonstrate conserved propofol binding sites involving the S4 voltage sensors and the S4-S5 linkers in NaChBac and NavMs, and also suggest state-dependent changes at these sites. Then, using molecular dynamics simulations to elucidate the structural basis of propofol modulation, we show that the S4 voltage sensors and the S4-S5 linkers shape two distinct propofol binding sites in a conformation-dependent manner. These interactions help explain how propofol binding promotes activation-coupled inactivation to inhibit Nav channel function.

## Introduction

General anesthetics comprise a class of chemically heterogeneous drugs capable of simultaneously inducing loss of consciousness, amnesia, immobilization, and analgesia through mechanisms that are not well understood (Hemmings, 2019). Propofol, one of the most effective and widely used intravenous general anesthetics, was invented in the late 1970’s and is used for the induction and maintenance of sedation and general anesthesia (Sahinovic, 2018; Walsh, 2018). Despite favorable properties, such as rapid induction and recovery times, propofol also has significant negative cardiopulmonary side effects, including apnea, hypotension, impaired myocardial contractility, and dystonia (Sahinovic, 2018). The hypnotic effects of propofol result primarily from direct interactions with postsynaptic GABA_A_ receptors, which potentiate inhibitory GABA activity in the brain (Eckenhoff, 2018). However, general anesthesia likely involves multiple targets that may be responsible for other endpoints and side effects (Eckenhoff, 2018; Hemmings, 2019). In particular, voltage-gated Na^+^ channels (Navs), which are responsible for the initiation and propagation of action potentials in the nervous system, may be one of these targets. Studies show that, general anesthetics typically inhibit Navs (Ouyang, 2003, 2007; Barber, 2014; Sand, 2017; Yang, 2018). A current hypothesis proposes that the negative modulation of Navs by general anesthetics may contribute to the presynaptic effects of these drugs on neurotransmission (Herold, 2012; Zhou, 2019). This negative modulation may result from various possible mechanisms in which anesthetic binding to Navs impairs ion permeation and/or modifies gating. Determining the biophysical and structural bases of these mechanisms is of paramount importance in the quest to understand the fundamental principles that explain how general anesthetics affect the function of diverse ion channels and thereby induce physiological effects (Covarrubias, 2015).

Recent work demonstrated that negative modulation of the prokaryotic Nav NaChBac by the inhalational anesthetics isoflurane and sevoflurane may result from direct interactions that promote both activation and inactivation (Ouyang, 2007; Barber, 2014; Kinde, 2016; Sand, 2017). We have also found that, in contrast to the mechanism of local anesthetic action, propofol does not act as a pore blocker of NaChBac and another prokaryotic Nav NavMs (Yang, 2018). Instead, propofol renders voltage-dependent activation more favorable, thereby promoting activation-coupled inactivation and, consequently, net negative modulation (Yang, 2018). Furthermore, molecular dynamics simulations of the prokaryotic Navs in the open/inactivated state revealed preferential propofol binding to an intersubunit hydrophobic pocket involving the S4-S5 linker (Yang, 2018), which was corroborated by NMR-based experiments in NaChBac (Wang, 2018). The S4-S5 linker plays a vital role in the electromechanical mechanisms of voltage-gated ion channels by providing a physical connection between the voltage sensing and pore domains, (Catterall, 2010; Wisedchaisri, 2019). Therefore, propofol may act as a gating modifier of Navs that promotes activation-coupled inactivation by interacting with the S4-S5 linker and neighboring regions. However, a conclusive demonstration of propofol binding sites in Navs is still lacking. Understanding the structural properties of these sites is a key step toward establishing the mechanism of propofol action.

To close this critical gap and render a structural model of propofol action, we performed unbiased photoaffinity labeling (PAL) experiments using AziP*m*, a pharmacologically active diazirine derivative of propofol (Hall, 2010). Results from NaChBac and NavMs show that propofol binds to critical pockets involving the S4 voltage sensor and the S4-S5 linker. Moreover, a distinct photolabeling pattern of a non-inactivating NaChBac mutant (T220A) suggests that these interactions are state-dependent. Then, using the latest structural data from both prokaryotic and eukaryotic Navs, we conducted MD simulation studies to produce a detailed atomistic model of how propofol modulates Navs by interacting with the electromechanical apparatus in the resting, open, and inactivated states. The structural and mechanistic insights provided by this study advance our understanding of how propofol modulates voltage-gated ion channels, which may contribute to endpoints and side effects of propofol-induced anesthesia.

## RESULTS

### Modulation of NaChBac gating by AziP*m*

To directly identify the propofol binding site in prokaryotic Navs, we employed an unbiased photoaffinity labeling (PAL) approach that has been extensively used to identify anesthetic binding sites of numerous proteins (Hall, 2010; Chiara, 2013; Weiser, 2013; Jayakar, 2014; Woll, 2016, 2017, 2018; Bensel, 2017). Here, we used AziP*m* (*m-*Azipropofol), an alkyl-diazirinyl derivative of propofol, to label propofol binding sites in NaChBac and NavMs (Hall, 2010). Though AziP*m* retains the pharmacological properties of propofol, we first sought to demonstrate that AziP*m* retains activity on prokaryotic Navs that is similar to that of the parent compound propofol.

Using whole-cell patch clamp electrophysiology, we characterized voltage-dependent gating of NaChBac heterologously expressed in HEK-293T cells, before and after exposure to 1 and 5 μM AziP*m* (**Figure 1A**, **Figure S1**). Previously, we reported that propofol hyperpolarizes the voltage dependencies of both activation and inactivation, and accelerates macroscopic inactivation (Yang, 2018). Here, we found that the modulations of NaChBac by AziP*m* and propofol are very similar (**Table 1**). AziP*m* induced parallel hyperpolarizing shifts of the peak conductance-voltage (G-V) relation (**Figure 1B**), without significant changes in the maximum conductance or effective gating charge of activation (**Figure 2A, middle and right**). At 1 and 5 μM AziP*m*, the ΔV_1/2_ of activation was −11.29 ± 1.44 mV and −12.34 ± 1.19 mV, respectively (**Figure 2A, left**). We also characterized the time constants of current decay to evaluate macroscopic inactivation and found that AziP*m* uniformly accelerates current decay between −40 and +60 mV in a concentration-dependent manner (**Figure 1C**). For example, the time constants of inactivation at +40 mV were 113 ± 12 and 54 ± 6 ms, in the presence of 1 and 5 μM AziP*m*, respectively, compared to 158 ± 12 ms for control (**Table 1**). AziP*m* also induced parallel hyperpolarizing shifts of the pre-pulse inactivation curves (**Figure 1D**), without significant changes in the effective gating charge of inactivation (**Figure 2B, right**). The ΔV_1/2_ of inactivation at 1 and 5 μM were −9.98 ± 0.98 mV and −18.95 ± 1.81 mV, respectively (**Figure 2B, left**). These hyperpolarizing shifts corresponded to 41 ± 2% and 47 ± 8% decreases in the channel availability at their respective baseline (control) V_1/2_ values of inactivation (**Figure 2B, right**).

**Table 1.**
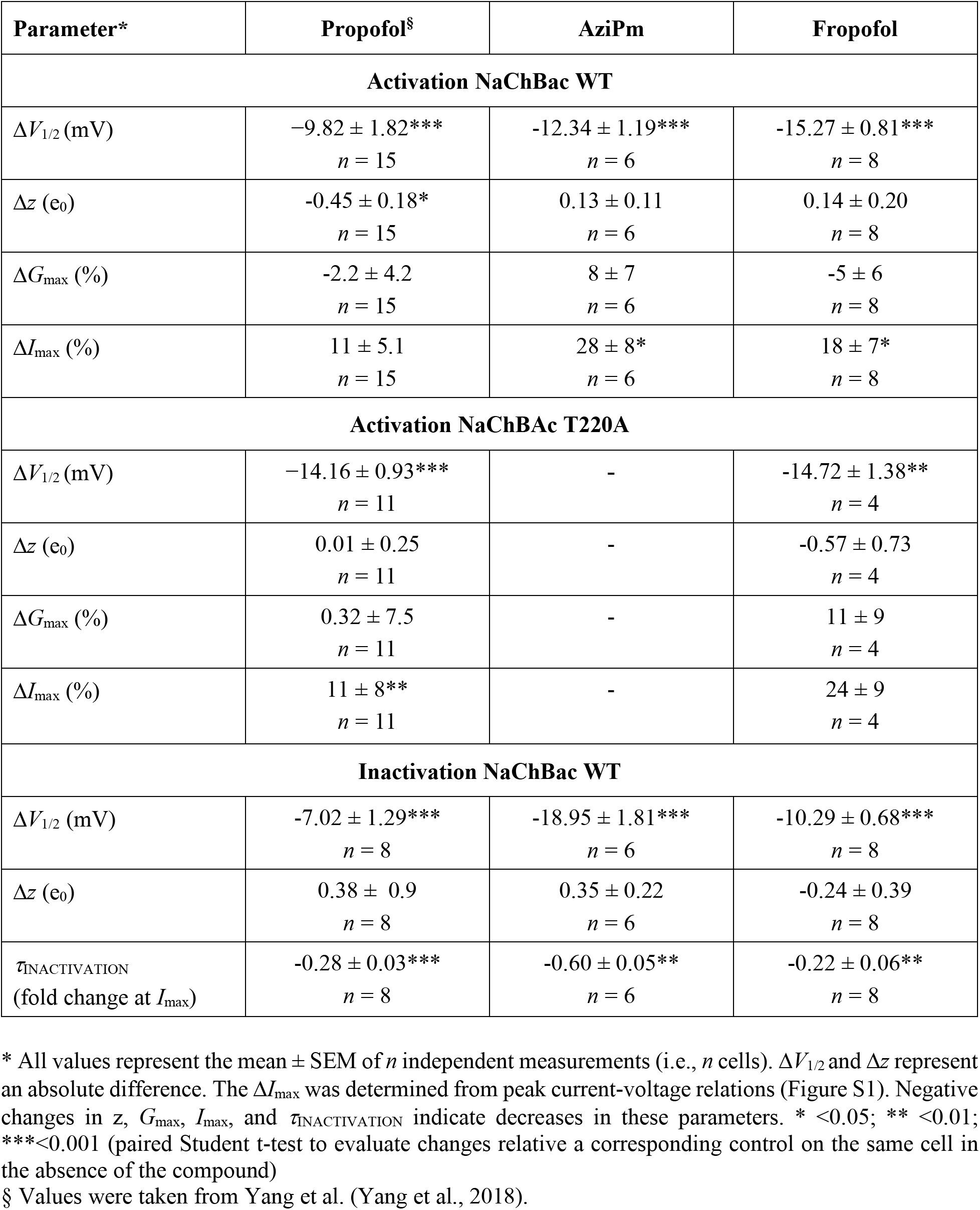
Changes in gating parameters of NaChBac induced by 5 μM propofol, AziP*m* and fropofol

| Parameter* | Propofol<sup>§</sup> | AziPm | Fropofol |
| --- | --- | --- | --- |
| **Activation NaChBac WT** |  |  |  |
| $\Delta V_{1/2}$ (mV) | $-9.82 \pm 1.82^{***}$<br>$n = 15$ | $-12.34 \pm 1.19^{***}$<br>$n = 6$ | $-15.27 \pm 0.81^{***}$<br>$n = 8$ |
| $\Delta z$ ($e_0$) | $-0.45 \pm 0.18^*$<br>$n = 15$ | $0.13 \pm 0.11$<br>$n = 6$ | $0.14 \pm 0.20$<br>$n = 8$ |
| $\Delta G_{\max}$ (%) | $-2.2 \pm 4.2$<br>$n = 15$ | $8 \pm 7$<br>$n = 6$ | $-5 \pm 6$<br>$n = 8$ |
| $\Delta I_{\max}$ (%) | $11 \pm 5.1$<br>$n = 15$ | $28 \pm 8^*$<br>$n = 6$ | $18 \pm 7^*$<br>$n = 8$ |
| **Activation NaChBac T220A** |  |  |  |
| $\Delta V_{1/2}$ (mV) | $-14.16 \pm 0.93^{***}$<br>$n = 11$ | - | $-14.72 \pm 1.38^{**}$<br>$n = 4$ |
| $\Delta z$ ($e_0$) | $0.01 \pm 0.25$<br>$n = 11$ | - | $-0.57 \pm 0.73$<br>$n = 4$ |
| $\Delta G_{\max}$ (%) | $0.32 \pm 7.5$<br>$n = 11$ | - | $11 \pm 9$<br>$n = 4$ |
| $\Delta I_{\max}$ (%) | $11 \pm 8^{**}$<br>$n = 11$ | - | $24 \pm 9$<br>$n = 4$ |
| **Inactivation NaChBac WT** |  |  |  |
| $\Delta V_{1/2}$ (mV) | $-7.02 \pm 1.29^{***}$<br>$n = 8$ | $-18.95 \pm 1.81^{***}$<br>$n = 6$ | $-10.29 \pm 0.68^{***}$<br>$n = 8$ |
| $\Delta z$ ($e_0$) | $0.38 \pm 0.9$<br>$n = 8$ | $0.35 \pm 0.22$<br>$n = 6$ | $-0.24 \pm 0.39$<br>$n = 8$ |
| $\tau_{\text{INACTIVATION}}$<br>(fold change at $I_{\max}$) | $-0.28 \pm 0.03^{***}$<br>$n = 8$ | $-0.60 \pm 0.05^{**}$<br>$n = 6$ | $-0.22 \pm 0.06^{**}$<br>$n = 8$ |

\* All values represent the mean ± SEM of $n$ independent measurements (i.e., $n$ cells). $\Delta V_{1/2}$ and $\Delta z$ represent an absolute difference. The $\Delta I_{\max}$ was determined from peak current-voltage relations (Figure S1). Negative changes in $z$ , $G_{\max}$ , $I_{\max}$ , and $\tau_{\text{INACTIVATION}}$ indicate decreases in these parameters. \* <0.05; \*\* <0.01; \*\*\*<0.001 (paired Student t-test to evaluate changes relative a corresponding control on the same cell in the absence of the compound)
<sup>§</sup> Values were taken from Yang et al. (Yang et al., 2018).

**Figure 1.**
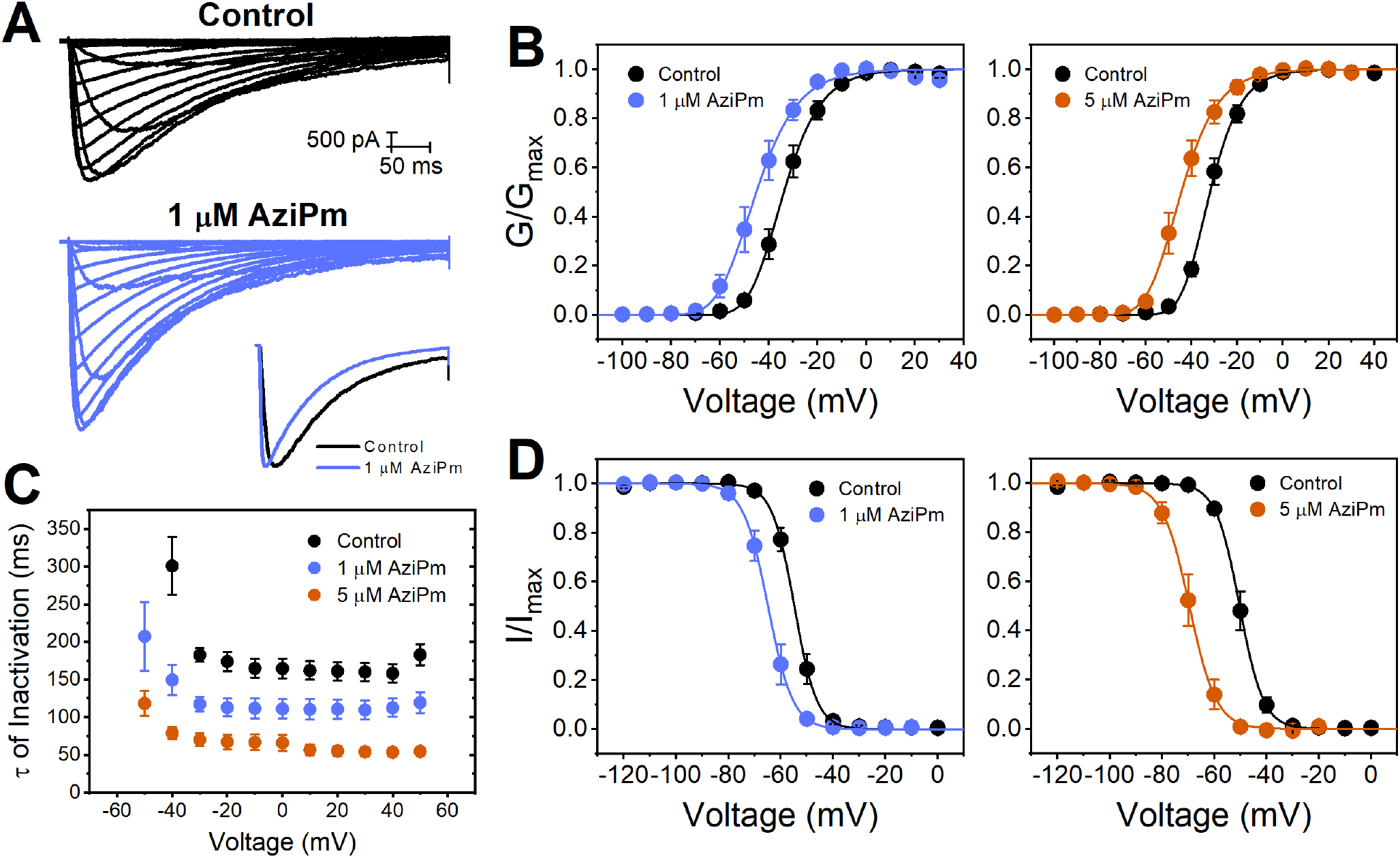
Modulation of NaChBac by AziP*m*. A) Families of whole-cell inward currents evoked by increasing step depolarizations from a holding voltage of −120 mV (−100 to +60 mV, ΔV = 10 mV). The start-to-start interval was 10 s. The inset shows scaled currents at 0 mV. B) Normalized peak conductance-voltage relations. The solid lines are best-fit 4^th^ order Boltzmann functions (summary of best-fit parameters on Table 1). C) Time constants of current decay against step depolarization voltage. The time constants were derived from fitting an exponential function to the decay of the current (Materials and Methods). D) Normalized pre-pulse inactivation curves. The currents were evoked by a constant step depolarization from increasing conditioning depolarizations as indicated on the graph The solid lines are best-fit 1^st^ order Boltzmann functions (summary of best-fit parameters on Table 1). NaChBac was expressed in HEK293T cells (Materials and Methods). **Source Data 1-5**.

**Figure 2.**
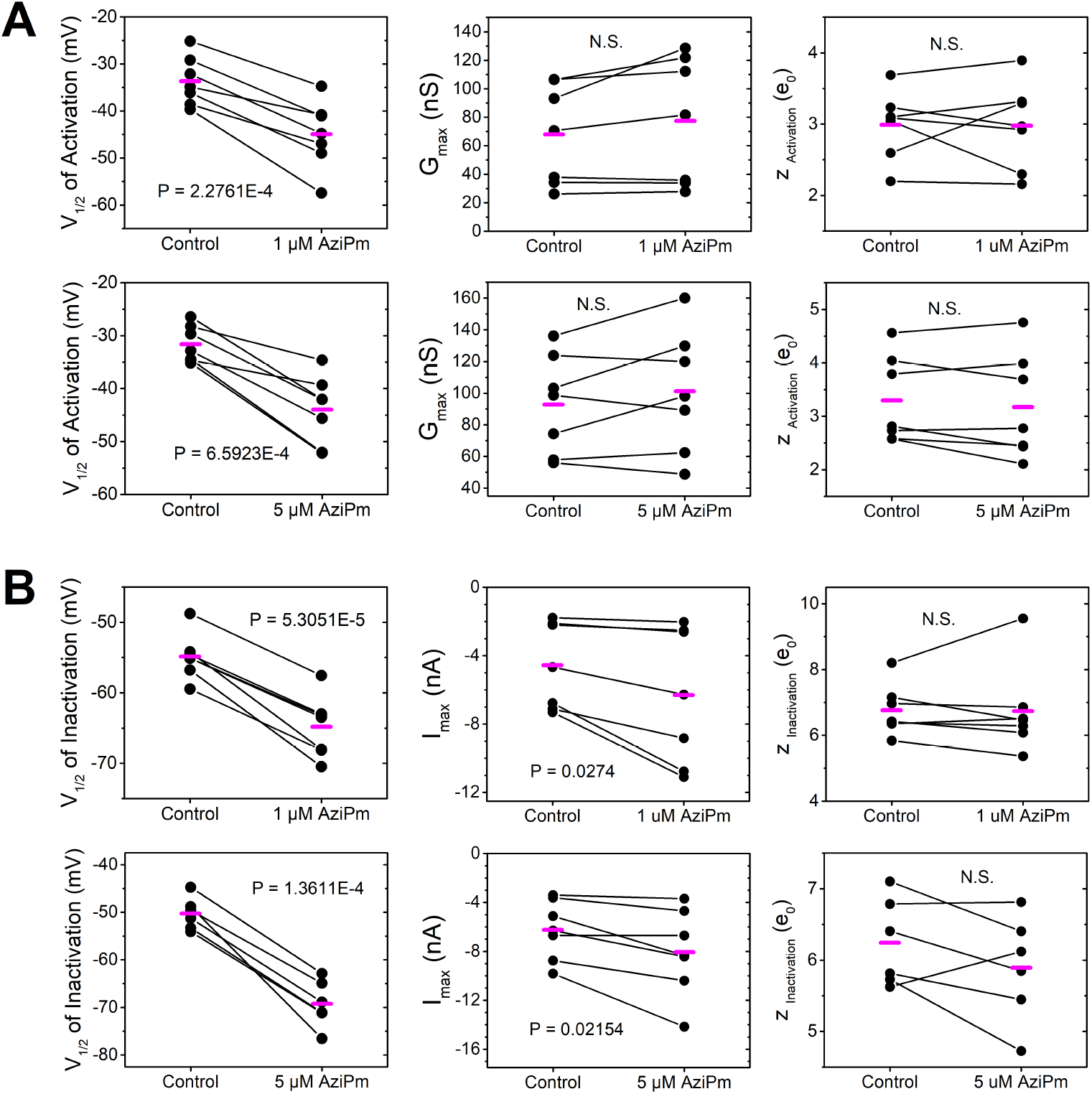
Statistical analysis of changes in NachBac gating parameters induced by 1 and 5 mM AziPm. A) and B) Scatter graphs of V_1/2_, G_max_ and z derived from the best-fit to 4^th^-order Boltzmann functions (Materials and Methods). C) and D) Scatter graphs of V_1/2_, I_max_ and z derived from the best-fit to 1^st^ order Boltzmann functions (Materials and Methods). The paired Student’s t-test was used to evaluate the changes. The p values are indicated in each graph (N.S. = not significant). **Source Data 1-5**.

Overall, these results demonstrate that AziP*m* induces a relative stabilization of both the open and inactivated states and accelerates macroscopic inactivation (**Table 1**). These changes closely resemble the modulation of NaChBac and NavMs by propofol and can be explained by the previously proposed mechanism (Yang, 2018). Like propofol, AziP*m* primarily promotes voltage-dependent activation and consequently favors activation-coupled inactivation, which is ultimately responsible for the net inhibitory action at steady state. Therefore, the similar modulatory behaviors of both propofol and AziP*m* reflects shared binding site(s) in prokaryotic Navs.

### AziP*m* binding to prokaryotic Nav channels

AziP*m* contains a trifluorodiazirine substitution at the *meta* position that generates a highly reactive and non-selective carbene moiety upon UV photoactivation, allowing the unbiased identification of propofol binding sites in NaChBac and NavMs. Using affinity-purified detergent-solubilized NaChBac (Materials and Methods; **Figure S2**), we found AziP*m* photomodification at I124, S125 and N146 (**Table 2**; **Figures S3a-d**). Whereas adducts at I124 and S125 were detected in separate trials, the adduct at N146 was detected once (**Table 2**; **Figures S3a-d**). The adducted residues span a region that includes the end of the S4 voltage sensor (I124 and S125), the S4-S5 linker and the beginning of S5 in the pore domain (N146; **Figures S6 and S7**). This is a region that undergoes critical conformational changes during activation and inactivation gating. Thus, to investigate a potential conformation dependence of propofol binding in NaChBac (Lee, 2012), we also performed photolabeling experiments on the non-inactivating NaChBac T220A mutant. Previously, we reported that propofol induces robust positive modulation of this mutant by stabilizing the open state in the absence of inactivation (Yang, 2018). We detected AziP*m* photomodification of NaChBac T220A at R131 at the N-terminal end of the S4-S5 linker, while neither I124 nor S125, which were labeled in WT NaChBac, were identified in separate attempts (**Table 2**; **Figures S4a-b**).

**Table 2.** AziP*m* Adduction of Prokaryotic Nav Channels in Multiple Independent Assays

| AziPm Adducted Peptide Fragments*<br>(bold letter highlights the adducted residue) |  |  |
| --- | --- | --- |
| NaChBac WT | NaChBac T220A | NavMs WT |
| <sup>120</sup>VLRA **I** SVVPSLR<sup>131</sup><br><sup>123</sup>A **I** SVVPSLR<sup>131</sup><br><sup>123</sup>AI **S** VVPSLRR<sup>132</sup><br><sup>135</sup>DALVMTIPALG **N** IL<sup>148</sup> | <sup>123</sup>AISVVPSL **R**<sup>131</sup><br><sup>123</sup>AISVVPSL **R**<sup>131</sup> | <sup>36</sup>**T** LSASSQNLLER<sup>47</sup><br><sup>110</sup>LVSV **I** PTMR<sup>118</sup><br><sup>110</sup>LVSVIPTM **R**<sup>118</sup> |

\* Each peptide sequence correspond to an independent experiment. The mass spectroscopy results and % sequence coverage from each individual attempt are shown under Supplemental Information (**Figures S3a-** **d, S4a-b, S5a-c**).

In NavMs, an AziP*m* photomodification was detected at R118, which is equivalent to R131 in NaChBac (**Table 2**; **Figure S5a-c**). I114 was also adducted in NavMs, a residue that is in close proximity to the equivalent NachBac residues, I124 and S125 (**Figure S6**). A sequence alignment that compares NaChBac and NavMs proteins demonstrates that the regional determinants of propofol binding are conserved in two distinct bacterial Nav channels. (**Figure S7**).

Consistent AziMP*m* photo-adduction limited to a discrete set of neighboring residues in two distinct bacterial Nav channels and a non-inactivating Nav mutant strongly suggests the presence of AziP*m* binding pockets mainly involving the S4 segment and the S4-S5 linker in NaChBac and NavMs. Moreover, the distinct photoadduction pattern of NaChBac T220A suggests that propofol binding in NaChBac depends on the gating conformation.

### MD simulations of Propofol binding to NaChBac in the inactivated state

Under the conditions of the photoloabeling experiments, NaChBac is most likely in the inactivated state (i.e., in the absence of a membrane potential). Thus, we characterized propofol binding to inactivated NaChBac in silico by conducting molecular dynamics (MD) simulations in the presence of an excess of propofol molecules to monitor binding near the experimentally observed sites, I124, S125, R131, and N146 (flooding MD simulations; Materials and Methods). Analysis of the four symmetry-related subunits of NaChBac consistently demonstrate two specific binding cavities for propofol: one located near the C-terminal end of S4 at the intracellular side of the voltage sensing domain, and the other, contiguous to the first, at the N-terminal side of the S4-S5 linker (**Figure 3A**). Whereas R131 and S125 line the first pocket and are in direct contact with bound propofol, N146 appears closer to the second pocket. I124 is in between both pockets and its sidechain points toward the membrane (**Figure 3B**). To ascertain that photoadduction events are possible between the bound propofol and sites I124, S125, R131, and N146, we calculated the minimum distance between these amino acids and the bound molecules of propofol for each configuration sampled along the simulated trajectory (we considered both main- and side-chain groups). The bimodal nature of the distribution of distances results from tumbling of the molecule within the binding pocket and suggests loose hydrophobic interactions between propofol and the pocket. Consistent with the photoadduction results, we found that propofol is within 4 Å from S125, R131, N146 in a significant fraction of the trajectory, which is 100 ns long (**Figure 3C**). By contrast, I124 appears to be farther away, suggesting that its interaction with propofol occurs in a different conformation (i.e. resting or open). Also, we observed no H-bonding between the 1-hydroxyl of propofol and groups in the channel protein, albeit a water molecule appears to bridge the propofol’s 1-hydroxyl to N146.

**Figure 3.**
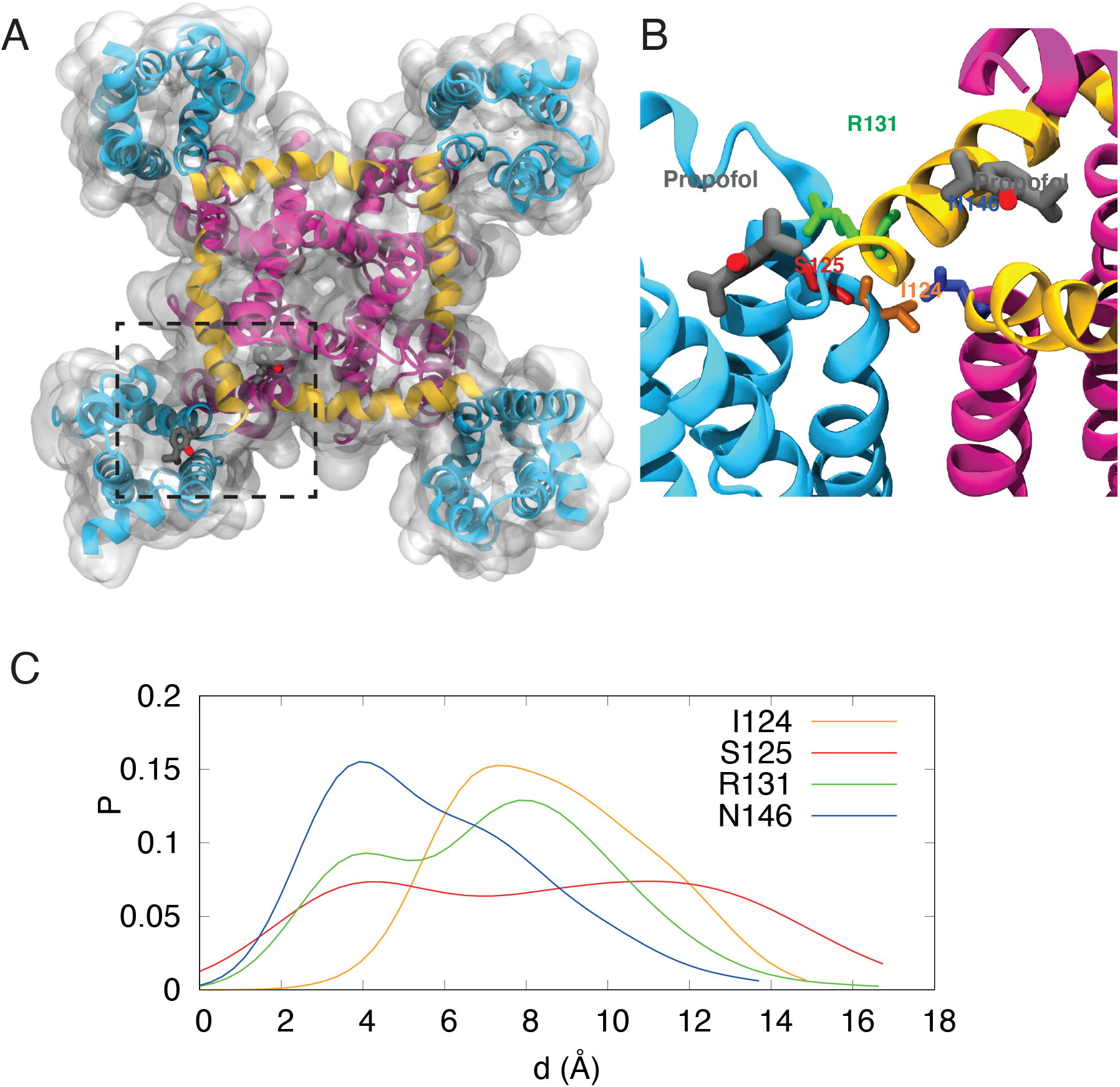
Binding of propofol to NaChBac in the inactivated state. A) Intracellular view of the tetrameric assembly. The voltage sensing (S1-S4) and pore domains (S5, S6) are colored in light blue and purple, respectively, and the S4-S5 linker connecting these domains is colored in yellow. Two propofol molecules are shown occupying two distinct pockets. One at the end of the S4 helix, and the other in close contact with the S4-S5 linker. B) Close up of the region highlighted in A), with amino acids I124 (orange), S125 (red), R131 (green) and N146 (blue) propofol (grey) shown as sticks. C) Minimum distance distribution between I124, S125, R131, N146 and propofol molecules.

### Propofol binding to NaChBac models in the resting, activated/open and inactivated states

To gain insights into the state-dependent character of the interaction between propofol and I124, S125, R131 and N146, we used comparative homology modeling to generate structural models for the resting and activated/open states (Materials and Methods). We first investigated whether or not the sidechains of the photoadducted residues face the pockets in any of the conformational states. Then, a comparative analysis of resting, activated/open, inactivated states focused on the size of the pockets allowed us to formulate hypotheses on the binding of propofol to distinct conformational states.

The major change concerns S125 and R131, which line the voltage sensor domain pocket in the activated/open and inactivated states, but not in the resting one (**Figure 4**). I124 is also particularly interesting because its sidechain faces the membrane in the inactivated and open states and is in close contact with N146 in the resting state, which may have a significant impact on photoadduction (**Figure 4**).

**Figure 4.**
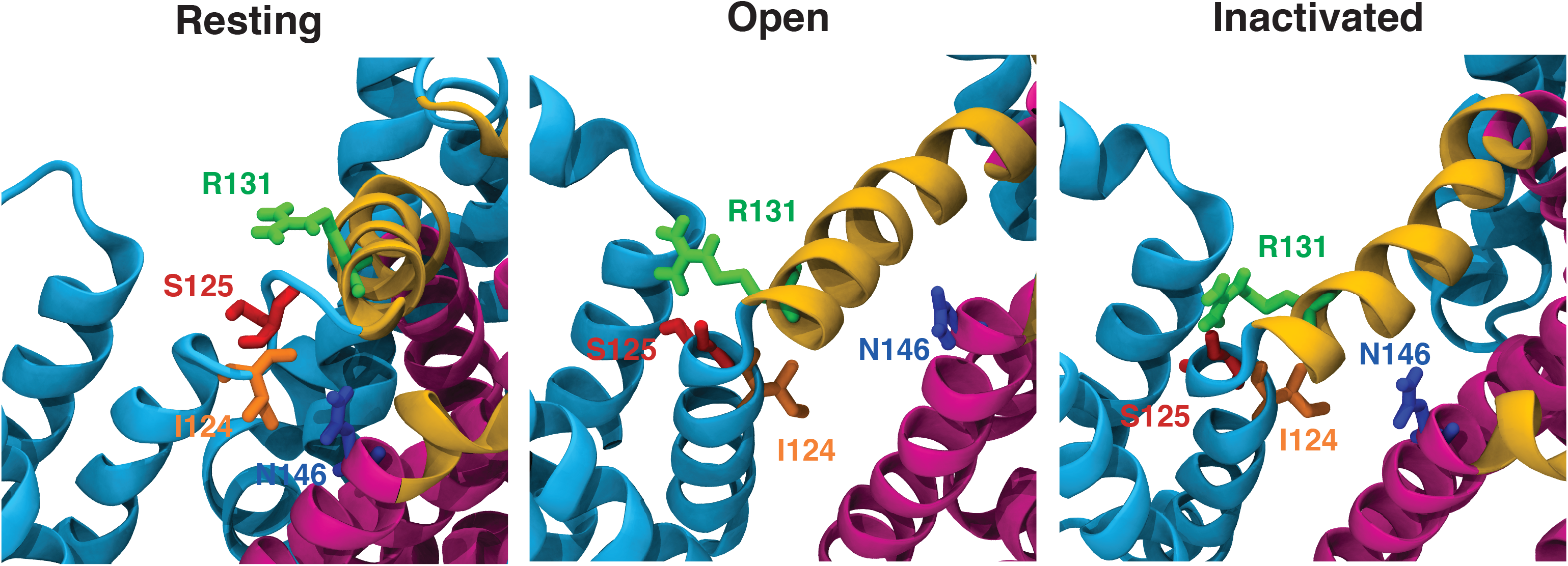
Models of the resting, open, inactivated states of NaChBac. Close ups of the propofol binding sites as described in Figure 3 legend. The voltage sensing (S1-S4) and pore domains (S5, S6) are colored in light blue and purple, respectively, and the S4-S5 linker connecting these domains is colored in yellow. The interacting amino acids are highlighted.

We then identified changes in the pocket geometry across the three states. The pocket facing the S4-S5 linker appears to be present only in the inactivated state (**Figure 5**). By measuring the number of waters molecules within 6 Å of residues L133 and I147 as a proxy for the pocket’s volume, we confirmed that this pocket is sufficiently large to bind propofol in the inactivated state only. By contrast, the site is collapsed in the resting and activated/open states (**Figure 5**). The pocket at the intracellular side of the voltage sensing domain is present in both the activated/open and inactivated states, while showing a complete reorganization in the resting one (as a result of the downward sliding of S4).

**Figure 5.**
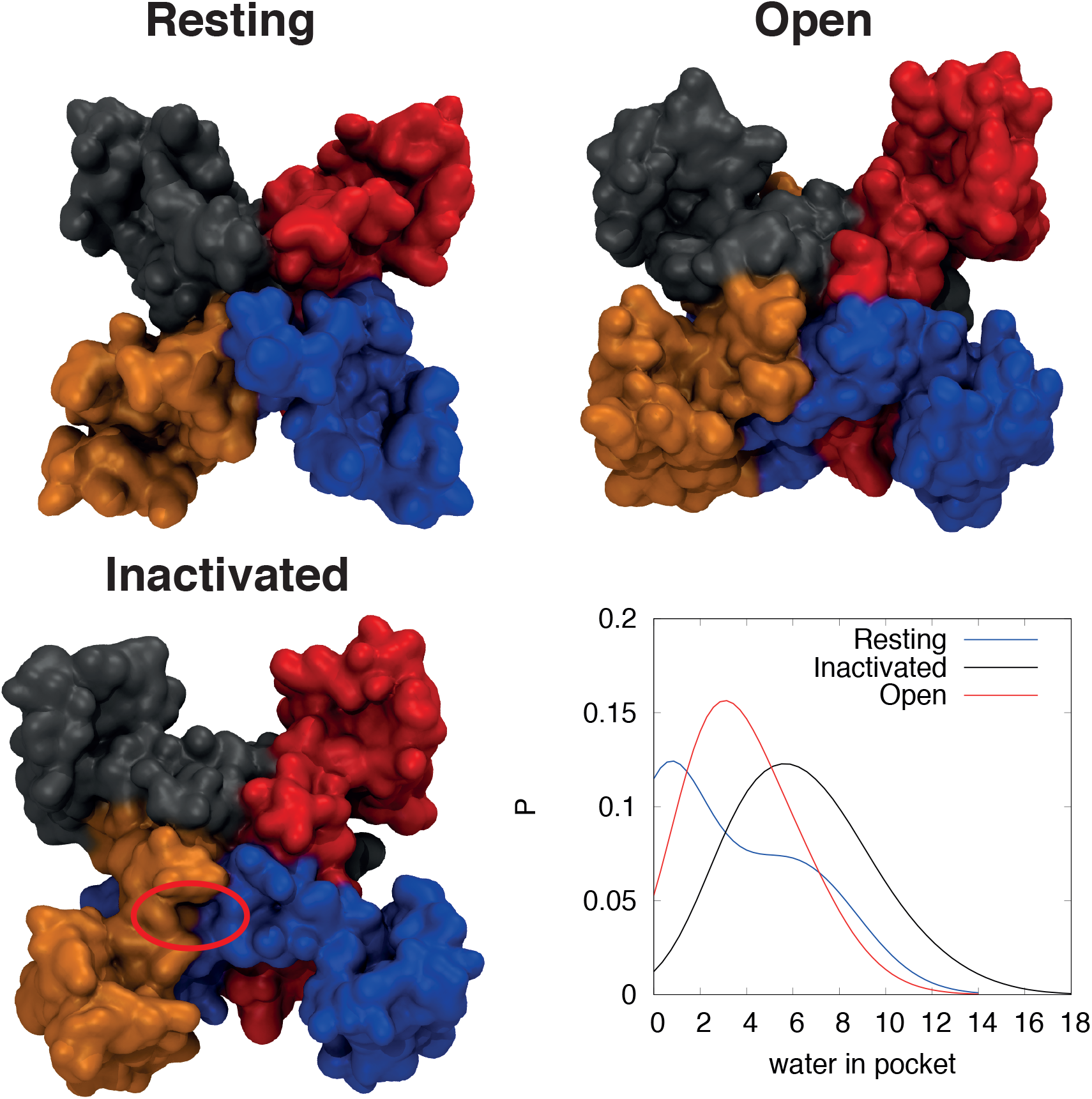
NaChBac models of resting, open, inactivated states. Molecular surfaces and colored according to the distinct symmetry related subunits. The S4-S5 linker pocket is highlighted in the inactivated state. Histograms show the number of water molecules that are closer than 6Å from amino acids L133 and I147.

### Structural conservation of the propofol binding site in NaChBac and NavMs

The electrophysiological and photoadduction experiments demonstrated extensive conservation in the ways in which propofol interacts with both NaChBac and NavMs (**Table 2**; **Figures S6 and S7**). To determine the structural correlates of this conservation, we compared the residues I124, S125, R131, N146 from NaChBac with the corresponding ones from the evolutionary related channel NavMs (PDB: 5HVX) (Sula et al., 2017). A structural superposition of the two structures in the open sate revealed that the equivalent NavMs residues are V111, S112, R118 and S133, respectively (**Figure 6**). This alignment confirms the importance of R118 (corresponding to R131 in NaChBac) as a determinant of propofol binding in NavMs. I114 is not among the four residues, albeit it was also phototadducted in NaMs; however, the peptide bond atoms of this residue are in close proximity to other residues that may line the voltage sensor site (**Figure 6**).

**Figure 6.**
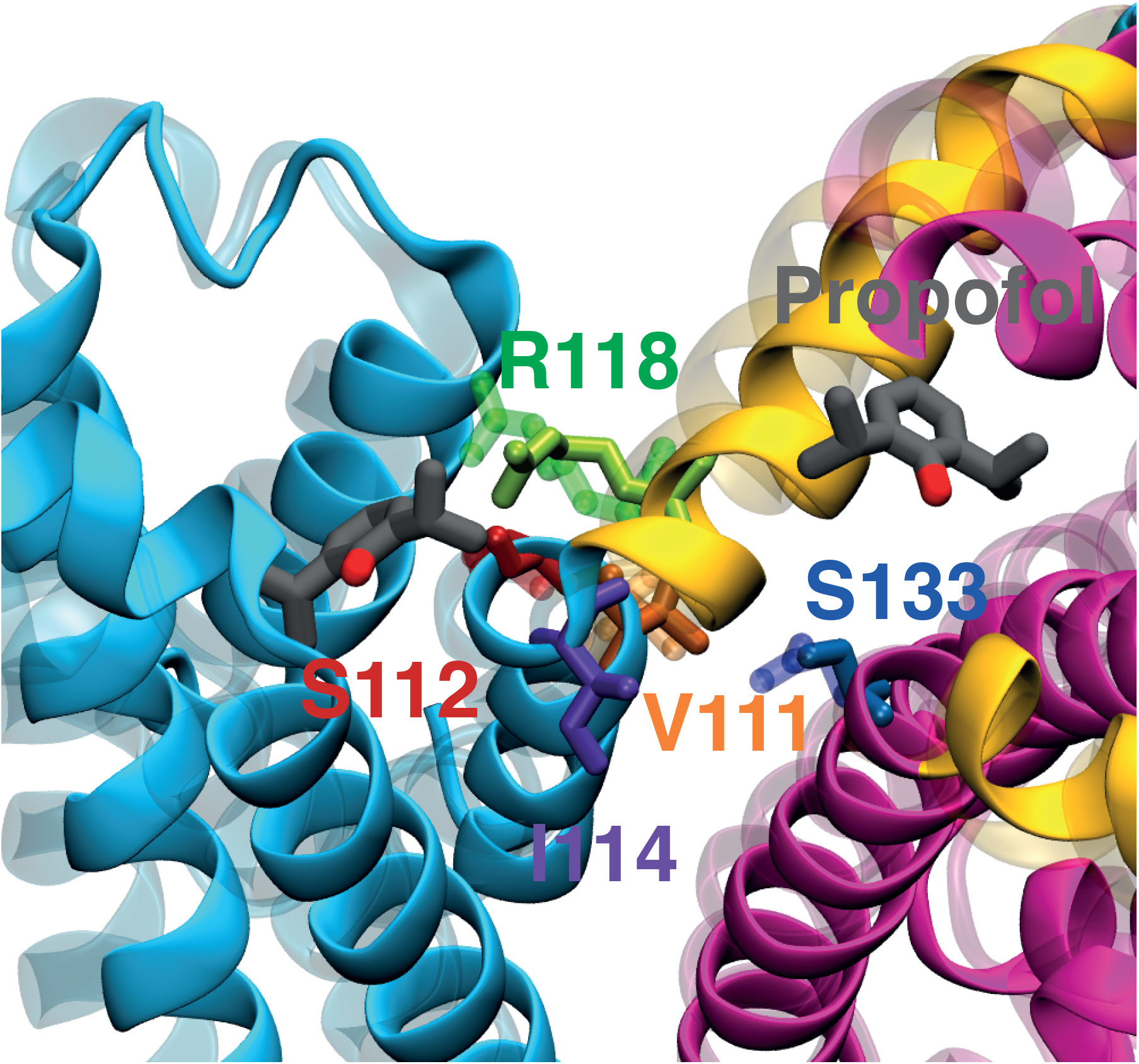
Structural superposition between NavMs and NaChBac. The amino acids aligned to I124, S125, R131, N146 are V111, S112, R118 and S133. I114 is also highlighted in violet.

### The hydroxyl group of propofol is not involved in NaChBac modulation

The MD simulations of propofol binding to NaChBac and NavMs did not reveal critical H-bond interactions involving the 1-hydroxyl group and protein groups. To experimentally test the role of propofol’s 1-hydroxyl group, we used fropofol, a propofol derivative in which the 1-hydroxyl group is replaced with an isosteric fluorine atom. Previously, we used fropofol to demonstrate that the propofol 1-hydroxyl group is necessary for the positive allosteric modulation of the ionotropic GABA_A_ receptor and the induction of hypnosis in tadpoles (Woll, 2015). Thus, we proposed that hydrogen bonding involving the 1-hydroxyl group is necessary for propofol binding in GABA_A_ receptors.

We found that, in ways that are reminiscent of propofol action, fropofol also modulates activation and inactivation gating (**Figure 7A-C; Table 1**). The time constants of current activation decreased gradually at all voltages between −40 and +40 mV (**Figure 7D**). At −40 mV, for example, the time constant of activation was 24 ± 3 and 7 ± 1 ms in the absence and presence of 5 μM fropofol, respectively. Although they do not exhibit significant voltage dependence between −40 and +40 mV, fropofol decreased the time constants of inactivation uniformly (**Figure 7E**). At +20, for instance, the time constant of inactivation was 122 ± 9 ms in the presence of 5 μM, compared to 168 ± 10 ms for the control condition. Fropofol also induced parallel hyperpolarizing shifts on the G-V relation and the pre-pulse inactivation curve. The ΔV_1/2_ of activation and inactivation induced by fropofol was −15.27 ± 0.81 mV and −10.29 ± 0.68 mV, respectively (**Figure 7A-C; Table 1**; **Figure S9**).

**Figure 7.**
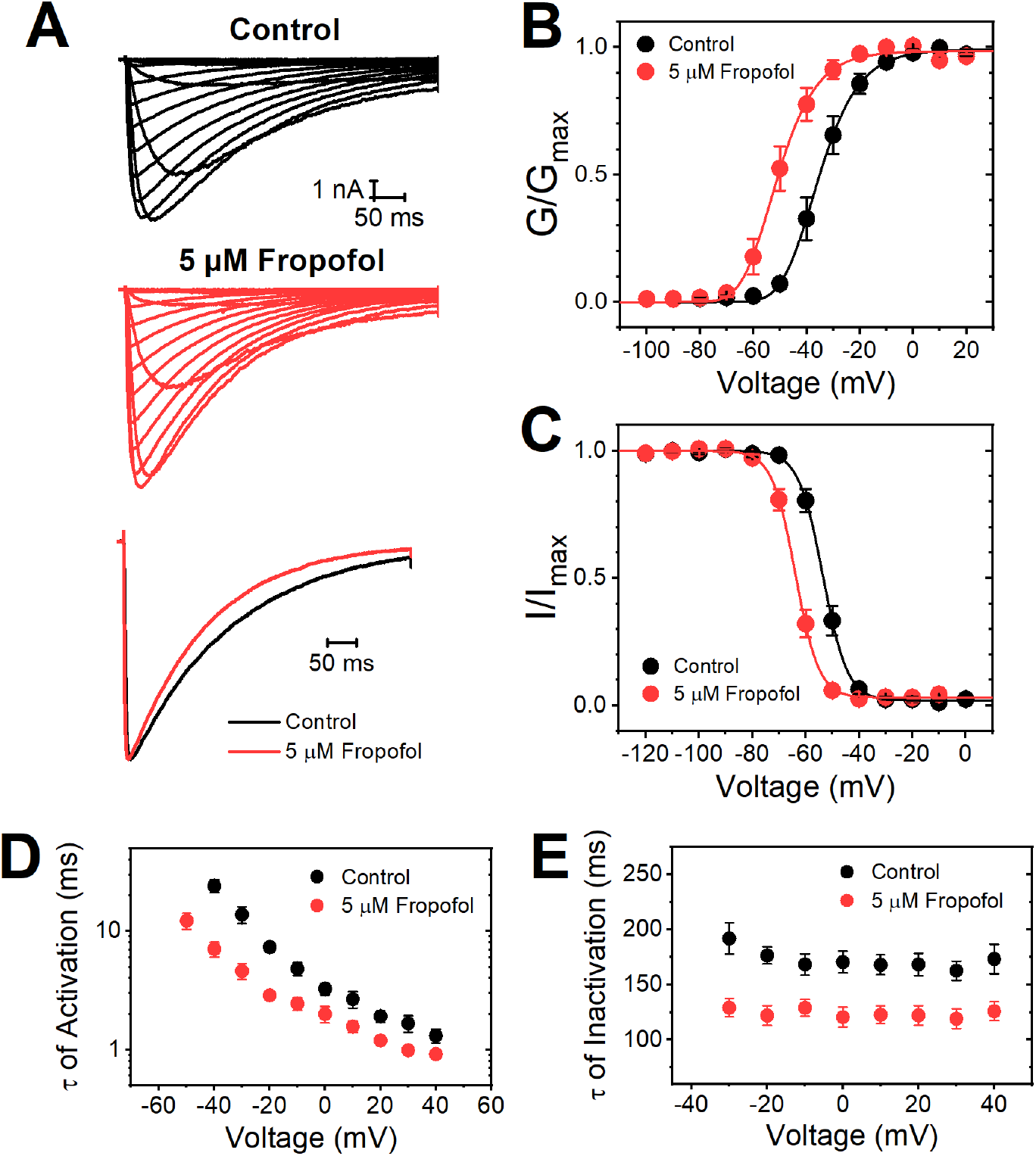
Modulation of NaChBac by fropofol. A) Families of whole-cell inward currents evoked by increasing step depolarizations from a holding voltage of −120 mV (−100 to +60 mV, ΔV = 10 mV). The start-to-start interval was 10 s. The overlay shows scaled currents at 0 mV. B) Normalized peak conductance-voltage relations. The solid lines are best-fit 4^th^ order Boltzmann functions (summary of best-fit parameters on Table 1). C) Normalized pre-pulse inactivation curves. The currents were evoked by a constant step depolarization from increasing conditioning depolarizations as indicated on the graph (Materials and Methods). The solid lines are best-fit Boltzmann functions (summary of best-fit parameters on Table 1). D) Time constants of current activation against step depolarization voltage. The time constants were derived from fitting an exponential function to the rising phase of the current (Materials and Methods). E) Time constants of current decay (inactivation) against step depolarization voltage. The time constants were derived from fitting an exponential function to the decay of the current (Materials and Methods). NaChBac was expressed in HEK293T cells (Materials and Methods). **Source data 6-8**.

These results demonstrate that the gating modulations of NaChBac by propofol and fropofol are overall similar. Providing further support to this conclusion, fropofol recapitulates propofol action by inducing a robust hyperpolarizing shift on the G-V relation of the non-inactivating mutant NaChBac T220A (**Figure S10**). The fropofol-induced ΔV_1/2_ of activation was −14.72 ± 1.28 mV (**Table 1**). These results strongly suggest that the 1-hydroxyl group of propofol is not involved in gating modulation of NaChBac by this anesthetic. Together with the AziP*m* photolabeling results, the electrophysiological characterizations of AziP*m* and fropofol actions provide sound experimental constraints on the generation of structural models to help explain propofol binding to prokaryotic Navs and the mechanism of gating modulation of these ion channels by propofol. Furthermore, these results shed light on potential clinical implications of the modulation of eukaryotic Navs by propofol.

## DISCUSSION

Previously, we demonstrated that propofol inhibits prokaryotic Navs by promoting activation-coupled inactivation (Yang et a., 2018), and further computational work and NMR-based characterization suggested that propofol mainly interacts with S4-S5 linker residues on the intracellular side of these Navs (Yang et al., 2018, Wang et al., 2018). Thus, we hypothesized that propofol binds to pockets formed by the S4-S5 linker and neighboring regions, and that this interaction is responsible for the net negative allosteric modulation of prokaryotic Navs. This is likely because the S4-S5 linker plays vital role in the gating mechanisms of voltage-gated ion channels (Catterall, 2010). However, direct evidence for propofol binding to the proposed pockets was lacking, and the possibility of gating-dependent conformational changes that affect propofol binding to these pockets had not been examined. To tackle these problems, we investigated NaChBac, NavMs, and a NaChBac mutant. Specifically, we used AziP*m*, a photoactivatable propofol analog, to conduct unbiased photoaffinity labeling experiments coupled to electrophysiological analysis and molecular dynamics simulations. The main results demonstrate that propofol binds to conserved NaChBac and NavMs pockets formed by an intracellular region that spans the end of the S4 voltage sensor, the S4-S5 linker and the beginning of the S5 pore segment. Furthermore, we provided biochemical and computational evidence to suggest that the binding pockets undergo gating-dependent conformational changes, and that the 1-hydroxyl group of propofol is not critical for propofol binding to prokaryotic Navs.

### The structural basis of the allosteric modulation of prokaryotic Navs by propofol

Based on electrophysiological analysis and kinetic modeling that assumes preferential closed-state recovery from inactivation, we previously suggested that propofol binding to NaChBac and NavMs accelerates the rate constants of activation and inactivation without a change in the kinetics of recovery from inactivation (Yang et al., 2018). Without a direct identification of the propofol binding site(s), however, the structural correlates of this allosteric modulation remained speculative. The new photoadduction data and congruent MD simulation results now provide physical evidence for two propofol binding pockets in a region that is directly involved in the voltage-dependent communication between the voltage sensing domain and the pore domain. Propofol interacts with the intracellular side the S4 voltage sensor in one pocket, and the S4-S5 linker in the other (Figs. 8-10). Suggesting a conformation-dependent interaction, the inactivating NaChBac wild type and the non-inactivating NaChBac T220A mutant yielded different photoadduction patterns. Whereas AziP*m* adducts were found at I124, S125 and N146 in the wild type, the adduct at R131 was only found in the non-inactivating mutant. R131 corresponds to R118 in NaMs, which was also photoadducted with AziP*m* (Table 2). This result was at first surprising because NavMs currents undergo fast and complete inactivation. There are, however, important differences between these prokaryotic Navs that must be considered to understand this result. Compared to NaChBac, NavMs exhibits faster activation and inactivation kinetics and operates over a voltage range that is 30 – 40 mV more hyperpolarized (Yang et al., 2018). Furthermore, their S4 and S4-S5 linkers are only ~35% identical, and the photoadducts may form at the peptide bonds rather than the amino acid sidechains. Given these considerations, subtle structural differences in the propofol pockets between NavMs and NaChBac may explain why equivalent photoadducts are observed in potentially distinct conformations. Consistent with this possibility, a second photoadduct was found at I114 in NavMs, a position that is only two residues downstream from a position equivalent to S125 in NaChBac. Although we observed no exact match between the AziP*m* photoadducts detected in NaChBac and NavMs, it is striking that the vast majority of them were found in the S4-S5 linker or in close proximity to it in 8/9 reactions (**Table 2**). We therefore conclude that the S4-S5 linker and neighboring regions are direct physical determinants of propofol binding in prokaryotic Navs. This conclusion is also supported by the results of a previous study that used ^19^F-NMR saturation transfer difference spectroscopy to investigate the binding of 4-fluoropropofol to potential sites suggested by docking in silico (Wang et al., 2018). This study, however, identified additional sites in the pore, the external apex of the voltage sensor and the internal activation gate, which were not observed in the PAL experiments reported here. Various factors could have contributed to this discrepancy. For instance, NMR experiments required cysteine mutations at predetermined putative sites to label the protein with the NMR probe. These mutations and the probe can have effects on gating and propofol binding. These experiments also required concentrations of the fluorinated ligand that are 40 – 200 times higher than the concentrations of AziP*m* used in PAL experiments and, therefore, the additional sites might be low affinity sites that were not detected in our experiments.

The general convergence of extensive electrophysiological analysis, consistent photoadduction results and MD simulations extended to major gating states, helps now formulate a structure-based model to explain the mechanism of the allosteric modulation of prokaryotic Navs by propofol. Primarily, binding of propofol to the intracellular voltage sensor pocket in the activated state may catalyze the outward movement of the S4 segment and thereby promote complete activation and pore opening. In turn, this positive modulation, could also indirectly favor inactivation of open channels, which may start in the selectivity filter and propagate to the rest of the pore domain (Pavlov et al., 2005; Gamal El-Din et al., 2019). Propofol may also have catalytic allosteric effects on inactivation gating through its binding to the S4-S5 linker pocket in a putative inactivated state. This interaction is especially intriguing because this pocket in NaChBac and NavMs corresponds to a hydrophobic cavity recently identified as the binding site of the IFM motif in the eukaryotic Nav1.4 and Nav1.5 (Pan et al., 2018; Jiang et al., 2020). This motif is the “inactivation particle” of most eukaryotic Nav channels, which acts allosterically on the pore domain to induce fast inactivation (Catterall, 2012; Ahern et al., 2015).

### Conservation of general anesthetic sites involving the S4-S5 linker of diverse ion channels

Using photoaffinity labeling, previous work from our laboratories have provided strong evidence to suggest that the S4-S5 linker and neighboring regions of Kv and TRPA1 channels are critical structural determinants of their modulation by volatile (sevoflurane) and intravenous (propofol) anesthetics (Bu et al., 2017; Woll et al., 2017). Based on the new results reported here, we can reach a similar conclusion for prokaryotic Navs. There is, however, little to no identity between the S4-S5 linkers of distantly related Kv, TRPA1 and Nav channels, albeit they all have similar topological features (6 transmembrane, 6TM, segments) and overall architecture. Therefore, we propose that common physical-chemical properties (e.g., amphipathic character) and a similar shape of the cavities formed by this linker and neighboring regions (the S6 segment and possibly membrane lipids) are sufficient to support a favorable interaction with volatile anesthetics and propofol. In this scenario, the S4-S5 linkers and neighboring regions of a subset of 6TM ion channels are binding determinants of general anesthetics and are directly involved in the mechanism of action.

### Mechanistic implications for eukaryotic Navs

Whether or not the hydrophobic IFM cavity also binds propofol in eukaryotic Navs needs to be determined. Previous studies have demonstrated that propofol and volatile anesthetics inhibit mammalian Navs, and have suggested that this inhibition may depress excitatory synaptic transmission in the brain, but the molecular mechanism of this inhibition is not known (Ratnakumari and Hemmings, 1997; Lingamaneni, Birch and Hemmings, 2001; Ouyang, Wang and Hemmings, 2003; Herold and Hemmings, 2012). The study of prokaryotic Navs with a simpler homotetrameric structure has helped us generate a sound working hypothesis that explains how propofol can modulate Nav channel activation and inactivation gating through allosteric interactions that involve the S4 voltage sensor and the S4-S5 linker. Although these regions have structural features that are generally conserved in all Navs, eukaryotic Navs are more complex pseudo-tetrameric proteins made of four homologous domains; therefore, characterizing the mechanism of propofol action and its structural basis in detail has been intrinsically difficult (Catterall, 2012). Recent advances, however, have made possible the structural characterization of eukaryotic Navs with atomic resolution, providing avenues to visualize their interactions with toxins, anesthetics and other drugs (Pan et al., 2018; Shen et al., 2019; Jiang et al., 2020).

### Nav channel modulation by propofol and the end points of general anesthesia

Negative modulation of Navs would inhibit excitation of neurons and muscles and could be linked to common endpoints of general anesthesia, such as loss of consciousness, amnesia, analgesia and immobilization. Most likely, however, propofol-induced sedation and loss of consciousness result from potentiation of GABA receptors (Tang and Eckenhoff, 2019). In support of this conclusion, the propofol derivative fropofol, which has the 1-hydroxyl group replaced with an isosteric fluorine, does not interact with the GABA_A_ receptor and has no hypnotic activity (Woll et al., 2018). It was thus proposed that the 1-hydroxyl group of propofol participates in hydrogen bonding that is critical for the hypnotic activity of propofol resulting from its interaction with GABA_A_ receptors. In contrast, we found that propofol and fropofol have a very similar profile of activity on the prokaryotic NaChBac, indicating that H-bonding mediated by the propofol’s hydroxyl group does not play a critical role in a prokaryotic Nav (**Figure 6**). We do not know whether this would also be the case in eukaryotic Navs; however, such a difference would suggest that the documented inhibition of mammalian Navs by propofol may rather be linked to other pharmacological properties (e.g., muscle relaxant and anti-epileptic) or toxicity associated with myocardial depression (Lundstrom et al., 2010; Woll et al., 2015).

### Conclusions

We used a set of complementary approaches including electrophysiological analysis, mutagenesis, unbiased PAL, MD simulations and a manipulation of propofol chemistry to identify the propofol binding sites in two prokaryotic Navs and determine a plausible mechanism of action. We conclude that propofol binds to two contiguous binding cavities. One involving the cytoplasmic side of the S4 voltage sensor and the other involving the S4-S5 linker and part of the S5 segment. State-dependent interactions with these sites are the basis of a mechanism in which propofol promotes activation-coupled inactivation to induce net inhibition of the Navs. Given that the interactions do not depend on the 1-hydorxyl group of propofol, we hypothesize that the modulation of eukaryotic Navs by propofol may not be involved in the hypnotic effects of this anesthetic.

## MATERIALS AND METHODS

### Molecular and Cell Biology

Wild-type (WT) NaChBac cDNA in a modified pTracer-CMV2 expression vector was a gift from D. Ren (University of Pennsylvania, Philadelphia, PA). cDNA was amplified in bacterial culture and purified with QIAGEN™ Plasmid Midi Kit. To generate NaChBac T220A, a point mutation was introduced into the WT plasmid using the QuikChange site-directed mutagenesis method (Agilent). HEK-293 cells were transiently transfected with NaChBac WT or T220A cDNA using Lipofectamine™ 2000 transfection reagent (Invitrogen) and seeded onto 12 mm circular glass coverslips 24 hours prior to patch clamp recording. Standard protocols were followed for growth and maintenance of cells in culture.

### Electrophysiology

Patch pipettes were pulled from borosilicate capillary glass (LA16, Dagan Corp.) with a HEKA PIP6 micropipette puller (HEKA). Prior to recording, patch pipettes were fire polished to a final resistance of 1.5 to 2.3 MΩ. Whole-cell patch clamp recording was performed using an Axopatch 200B amplifier (Molecular Devices) and Digidata 1440A analog-to-digital converter (Molecular Devices). Series resistance was compensated at least 85%. Passive leak current and capacitive transients were subtracted online by standard P/4 protocol. All recordings were low-pass Bessel-filtered at 2 kHz and digitized at 15.4 kHz. Clampex 10 (pCLAMP 10, Molecular Devices) was used to control voltage protocols and for data acquisition.

The extracellular bath solution contained (in mM) 140 NaCl, 4 KCl, 1.5 CaCl2, 1.5 MgCl2, 10 HEPES, and 5 D-glucose, pH 7.3 adjusted with NaOH; the intracellular pipette solution contained (in mM) 15 NaCl, 80 CsF, 40 CsCl, 10 EGTA, and 10 HEPES, pH 7.3 adjusted with CsOH. AziP*m* was synthesized by previously published methods (Hall, 2010). Prior to use, AziP*m* was diluted in bath solution to working concentrations, followed by alternating sonication and vortexing for 4.5 minutes. All dilutions were prepared and utilized the same day. Cells were continuously perfused with bath solution at room temperature (22-25 °C) during recordings. In all experiments, all control recordings were collected first, prior to any AziP*m* exposure. Following control recordings, AziP*m* was perfused for ~3 minutes before collecting paired anesthetic recordings and continuously thereafter. To prevent inaccuracies due to any membrane lipid retention of anesthetic molecules and cumulative effects, each experimental cell was only exposed to a single concentration of AziP*m*, and washout data was not used.

### Voltage Protocols

In NaChBac WT and T220A, voltage-dependent activation was assessed with Na+ currents evoked by 500-ms depolarizing steps (−100 to +60 mV, ΔV = 10 mV) from a holding potential of −120 mV. Pre-pulse inactivation was assessed with a two-pulse protocol: (1) a 2-s conditioning pulse (−120 mV to −10 mV, ΔV = 10 mV), followed immediately by (2) a 50-ms test pulse to +10 mV. The holding potential was −120 mV.

### Analysis of Electrophysiological Results

Clampfit 10 (pCLAMP 10, Molecular Devices), Origin 9.1 (OriginLab Corp.) and Excel 2013 (Microsoft Corp.) were used to analyze voltage-clamp data. All evaluated parameters are reported as mean ± SEM. The paired samples t-test was used to assess differences between paired data sets in the absence and presence of propofol. P-values less than 0.05 are explicitly reported in the figures and figure legends, and “N.S.” indicates P ≥ 0.05.

Peak chord conductance (G) was calculated using *G* = *I* / [*V - V*_*rev*_], where *I* is the measured peak current, *V* is the command potential, and *V*_*rev*_ is the reversal potential extrapolated from individual current-voltage curves. The voltage dependence of activation (G-V curve) was derived from the best-fit fourth order Boltzmann function *G(V)* = [*G*_*max*_ / (*1 + e*^(*Vs – V)/k*^)]^4^ and normalized to *G*_*max*_, where *G*_*max*_ is the maximum peak conductance, *V*_*s*_ is the midpoint of activation for a single subunit, *V* is the command potential and *k* is the slope factor. The midpoint voltage of activation was calculated using *V*_1/2_ = (*V*_*s*_ + *1.67k)*. Pre-pulse inactivation parameters were determined from the best-fit first order Boltzmann function *I(V)* = *I*_*max*_ / (*1 + e*^(*V1/2 – V)/k*^) and normalized to *I*_*max*_, where *I*_*max*_ is the maximum current amplitude and *V*_*1/2*_ is the midpoint voltage of inactivation. Time constants of activation and inactivation were derived from the rising and decaying components of the Na^+^ current, respectively, using the best-fit single exponential of the form *I(t)* = *(A e*^*−t/τ*^ + *C)*, where *A* is the amplitude, *C* is the plateau constant, *t* is time, and *τ* is the time constant of activation or inactivation.

### Modeling and MD Simulations

The structural model of the open state was obtained using comparative homology modeling based on the cryoEM structure of the eukaryotic voltage-gated sodium selective channel NavPas (PDB: 6a90; Shen, et al., 2018). The inactivated conformation was modelled using NavAb as template (PDB: 5vb2; Lenaeus, et al., 2017). For the resting state we used the recently published xray structures of NavAb in the resting state (PDB: 6p6w; Wisedchaisri, et al., 2019). We threaded the sequence of NaChBac through the three PDB structures using the SWISS-MODEL software [Waterhouse, et al., 2018). The accuracy of the NaChBac model in the inactivated state was confirmed by comparing it against the recently published cryoEM structure of potentially inactivated NaChBac (**Figure S8**) (Gao et al., 2020).

Molecular systems were assembled using the CHARMM-GUI webserver (Wu, et al., 2014). Each channel state was embedded in a lipid bilayer of POPC, and the number of ions in the bulk was set to 0.15 M KCl and to obtain electrical neutrality. The molecular systems, included waters, contains about 180000 atoms. Molecular dynamics calculations are performed using the NAMD computational code (Phillips et al., et al., 2005), using the all-atom potential energy function CHARMM36 for protein and phospholipids, and the TIP3P potential for water molecules (Mackerell, et al., 2004; Jorgensen, et al., 1983). Periodic boundaries conditions are applied, and long-range electrostatic interactions are treated by the particle mesh Ewald algorithm (Essmann, et al., 1995). The Propofol molecule was modeled using the CGenFF webserver (Vanommeslaeghe, et al., 2012a; Vanommeslaeghe, et al., 2012b). Molecular systems are equilibrated for about 1 ns with decreasing harmonic restraints applied to the protein atoms and the lipid membrane. All trajectories are generated with a time step of 2 fs at constant normal pressure (1 atm) controlled by a Langevin piston and constant temperature (300 K) using a Nosé-Hoover thermostat. The configurations and results presented in Fig. 4D and Fig. 5 were obtained after simulating the three systems for about 100ns. Measurements related to Fig. 3A-C were obtained using flooding simulations based on the same protocol used previously (Wang, et al., 2018).

### Expression and Purification of NavMs and NaChBac

The genes for NaChBac and NavMs were amplified using appropriate PCR primers (**Table S1**) and subcloned into the in-house plasmid pETHT. This vector encodes an N-terminal His_6_ tag followed by a TEV protease recognition sequence, which is followed immediately by the sequence of the protein to be expressed. Both genes were inserted between the BsaI and NotI sites of the vector. The T220A mutant of NaChBac was prepared using PCR-mediated site-directed mutagenesis, with the primers shown in **Table S1**.

The NavMs plasmid was transformed into E. coli C41 (DE3). A single colony was diluted in LB medium containing 100 ug/mL ampicillin and grown overnight. The following morning, 20 mL of overnight culture were diluted into 1L of LB + ampicillin, and the flasks were shaken at 37°C until OD_600_ of 0.6-0.8, at which point IPTG was added at a final concentration of 0.5 mM. Cells were harvested 3.5 h after induction and frozen at −80° until use.

All purification steps were carried out at 4 °C. Cell pellets were thawed, re-suspended in lysis buffer (20 mM Tris-Cl, pH 7.5, 300 mM NaCl, 10 mM imidazole), and lysed by three passes through an ice-cold Avestin Emusiflex C5 cell disrupter. Cell lysates were clarified by centrifugation at 15,000×*g* for 15 mins; the cleared lysate was then centrifuged at 145,000x*g* for 1 hr. The resulting pellet was homogenized in lysis buffer, after which dodecyl maltoside was added to a final concentration of 1% (w/v). The solution stirred gently at 4 °C for 1 hr then centrifuged at 145,000 × *g* for 1 hr. The supernatant was filtered using a Millex-SV 5.0 um syringe filter (Merck-Millipore) and loaded onto a 1-mL HiTrap IMAC-HP column (GE Healthcare), which was pre-equilibrated with lysis buffer containing 0.1% dodecyl maltoside. The column was washed and then eluted with 50 mM Tris-Cl, pH 7.7, 500 mM NaCl, 500 mM imidazole, 0.1% dodecyl maltoside. The purified protein was then dialyzed against 50 mM Tris, pH 7.7, 500 mM NaCl, 0.1% DDM, and concentrated to a final concentration of 2-5 mg/mL (**Figure S2**).

NaChBac was expressed and purified in the same manner as NavMs, with the following modifications. After dilution of the original overnight culture, the flasks were shaken at 30 °C until an OD_600_ of 0.6-0.8 was reached, at which point IPTG was added and the temperature was reduced to 18 °C; cells were harvested 24 h after induction; cell pellets were disrupted in a lysis buffer containing 50 mM Tris, pH 7.7, 500 mM NaCl, 0.2 mM PMSF, 2 mg/mL lysozyme, 2 μg/mL DNase; and the detergent solubilization step was carried out overnight (**Figure S2**).

### Photolabeling of NaChBac WT, NaChBac T220A, and NavMs

A final concentration of 5 μM AziP*m* with or without 200 μM propofol was added to the purified NaChBac, NaChBac T220A, and NavMs to a final protein concentration of 1 μg/μl. The samples were equilibrated on ice in the dark for 5 min and then irradiated for 25 min at 350 nm with a RPR-3000 Rayonet lamp in 1-mm path length quartz cuvettes through a 295-nm glass filter (Newport Corporation).

### In-Solution Protein Digestion

After UV exposure proteins were precipitated overnight at −20 °C in 4 volumes of chilled acetone. Protein was pelleted for 20 min at 16,000 ×g at 4 °C then gently washed twice with 300 μl of chilled acetone. Protein pellets were air-dried before resuspension in 50 μl of 50 mM Tris-HCl, pH 8.0, 1% Triton X-100, and 0.5% SDS. Insoluble debris was pelleted by centrifugation at 16,000 × g. The samples were resuspension in final concentration of 50 mM NH_4_HCO_3_. Following 1 μL 0.5 M dithiothreitol (DTT) was added and samples were incubated at 56 °C for 30 min. 0.55 M iodoacetamide (IAA) was then added and protein samples were incubated at room temperature in the dark for 45 min. Sequencing grade-modified trypsin (Promega) was added to a final 1:20 protease: protein ratio (w:w) with additional of 0.2% (w/v%) ProteaseMax ™ Surfactant. Proteins were digested overnight at 37°C. Trypsin digested peptides were diluted to 200 μL with final concentration of 100 mM NH_4_HCO_3_ and 0.02% ProteaseMAX Surfactant prior to the addition of sequencing grade chymotrypsin (Promega) to a final 1:20 protease:protein ratio (w:w). Proteins were digested overnight at 37°C. Acetic acid (AcOH) was added to until the pH < 2 and the peptide digests were incubated at room temperature for 10 min prior to centrifugation at 16, 000 × g for 20 min to remove insoluble debris. The sample was desalted using C18 stage tips prepared in house. Samples were dried by speedvac and resuspended in 0.1% formic acid immediately prior to mass spectrometry analysis.

### In-Gel Protein Digestion

Photolabeled proteins were separated by SDS-PAGE. The identified bands corresponding to NaChBac and NavMs were excised. Excised bands were distained, dehydrated and dried by speed vac before proteins were reduced by incubation at 56° C for 30 min in 5 mM DTT and 50 mM NH_4_HCO_3_. The DTT solution was removed and proteins were then alkylated by the addition of 55 mM IAA in 50 mM NH_4_HCO_3_ and incubation at room temperature for 45 min in the dark. Bands were dehydrated and dried by speed vac before resuspension in 100 μL 0.2 % ProteaseMAX™ surfactant(Promega) and 50 mM NH_4_HCO_3_ solution containing trypsin (sequencing grade, Promega) at a 1:20 protease:protein ratio (w:w). Proteins were digested for 12-16 hrs at 37°C. After trypsin digestion, the samples were diluted to 200 μL with final concentration of 100 mM NH_4_HCO_3_ and 0.02% ProteaseMAX™ Surfactant. The samples were further digested overnight at 37 °C with the addition of sequencing grade chymotrypsin (Promega) to a final protease:protein (w:w) ratio of 1:20. Multiple peptide extractions were performed in order to increase hydrophobic peptide retrieval from the gel. The peptide extractions were pooled from the different ratio of acetylnitrile and acetic acid. The final extractions were sonicated for 20 min and dried by speed vac before resuspension in 0.5% acetic acid and further acidified until the pH < 2. Samples were sonicated for 10 min prior to centrifugation at 16, 000 x g for 20 min to remove insoluble debris. Samples were desalted using C18 stage tips prepared in house. Samples were dried by speed-vac and resuspended in 0.1% formic acid immediately prior to mass spectrometry analysis.

### Mass Spectrometry

Mass analysis was performed similar to as previously reported (Hall, 2010; Woll, 2017). Briefly, desalted peptides were injected into a Thermo LTQ Orbitrap XL Mass Spectrometer (Thermo Fisher Scientific, Waltham, MA, USA) or an Orbitrap Elite™ Hybrid Ion Trap mass spectrometer. Peptides were eluted with 100 min with linear gradients of ACN in 0.1% formic acid in water (v/v%) starting from 2% to 40% (85 min), then 40% to 85% (5 min) and finally 85% (10 min).

Spectral analysis was conducted using Maxquant (Thermo Scientific) to search b and y ions against the sequence containing NaChBac and NavMs. All analyses included dynamic oxidation of methionine (+15.9949 m/z) as well as static alkylation of cysteine (+57.0215 m/z; iodoacetamide alkylation). Filter parameters were Xcorr scores (+1 ion) 1.5, (+2 ion) 2.0, (+3 ion) 2.5, deltaCn 0.08, and peptide probability 0.05. Photolabeled peptides were searched with the additional dynamic AziP*m* modifications. Both the in-solution and in-gel sequential trypsin/chymotrypsin digests were searched without enzyme specification with a false discovery rate of 0.01. Samples were conducted in triplicate and samples containing no photoaffinity ligand were treated similarly to control for false positive detection of photoaffinity ligand modifications. To confirm the photolabeled adduct, MS work was repeated at the Proteomics Core Facility of the Wistar Institute (University of Pennsylvania, Philadelphia, PA, USA).

## ACKNOWLEDGMENT

This work was supported by following grants from the National Institutes of Health, USA: P01GM55876 (MC, VC, RGE), 1R01NS111997-01A1 (BAG), F30GM123612 (EY). We also thank members of the Covarrubias lab for their constructive feedback and support.

