## Supplemental and Source for "The propofol binding sites of prokaryotic voltage-gated sodium channels"

**CONTENT**

**Table S1. Primers used for NavMs and NaChBac sub-cloning and mutagenesis**

**Figure S1. NaChBac peak current-voltage relations in the absence and presence of AziP*m* or fropofol**

**Figure S2. Coomassie-stained SDS PAGE gels for purified NaChBac and NavMs proteins**

**Figure S3a. Mass spectra of AziP*m*-photolabeled peptides in NaChBac WT**

**Figure S3b. Mass spectra of AziP*m*-photolabeled peptides in NaChBac WT**

**Figure S3c. Mass spectra of AziP*m*-photolabeled peptides in NaChBac WT**

**Figure S3d. Mass spectra of AziP*m*-photolabeled peptides in NaChBac WT**

**Figure S4a. Mass spectra of photolabeled peptides in NaChBac T220A**

**Figure S4b. Mass spectra of photolabeled peptides in NaChBac T220A**

**Figure S5a. Mass spectra of photolabeled peptides in NavMs**

**Figure S5b. Mass spectra of photolabeled peptides in NavMs**

**Figure S5c. Mass spectra of photolabeled peptides in NavMs**

**Figure S6. Sequence coverage map for NaChBac, NaChBac T220A, and NavMs mass spectrometry analysis**

**Figure S7. Sequence Alignments of NaChBac and NavMs**

**Figure S8. Overlay of model and cryoEM structures of NaChBac**

**Figure S9. Statistical analysis of changes in NachBac gating parameters induced by 5 μM fropofol**

**Figure S10. Modulation of NaChBac T220A by fropofol**

**Table S1. Primers used for NavMs and NaChBac sub-cloning and mutagenesis**

| **NavMS sub-cloning primers** | |
| --- | --- |
| Forward | TTTTTTGGTCTCTCATGTCACGCAAAATCCGCGATTT |
| Reverse | TTTTTTAAGCTTATTATTTTTTTTGAGGTTGTTGACGTTG |
| **NaChBac WT sub-cloning primers** | |
| Forward | TTTTTTGGTCTCTCATGAAAATGGAAGCTAGACAGAAACAG |
| Reverse | TTTTTTGCGGCCGCACTATTATTTCGATTGTTTAAGCAAGCTTTTTAG |
| **NaChBac T220A mutagenesis primers** | |
| Forward | GTCTTAATCGGTGCGTTTATCATCT |
| Reverse | AGATGATAAACGCACCGATTAAGAC |

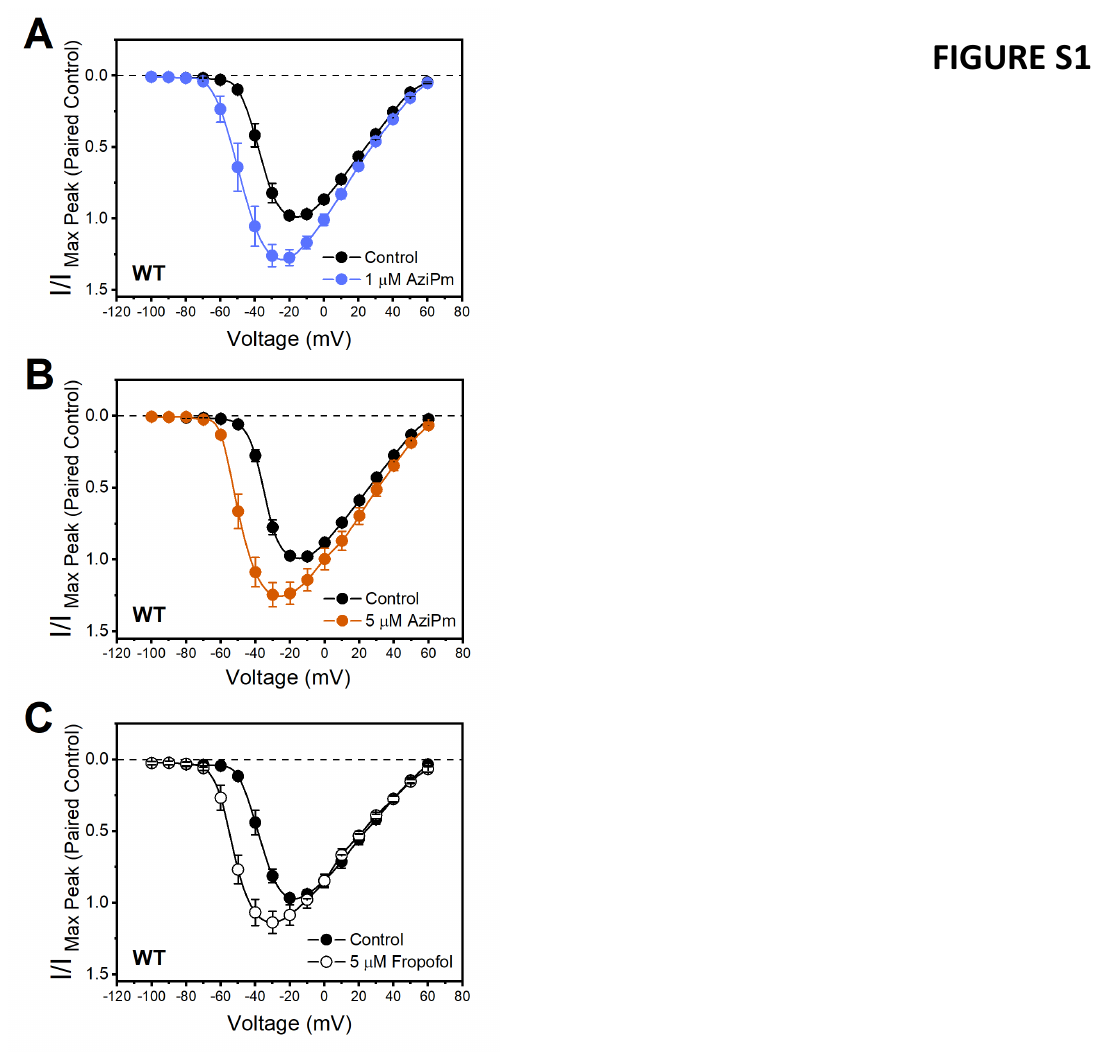

**Figure S1. NaChBac peak current-voltage relations in the absence and presence of AziP*m* or fropofol.** A) The effect of 1 mM AziPm. B) The effect of 5 mM AziPm. C) The effect of 5 mM fropofol. These relations are normalized relative to the corresponding control. The lines connecting the symbols are spline interpolations. Error bars indicate ± SEM. **Source Data 10.**

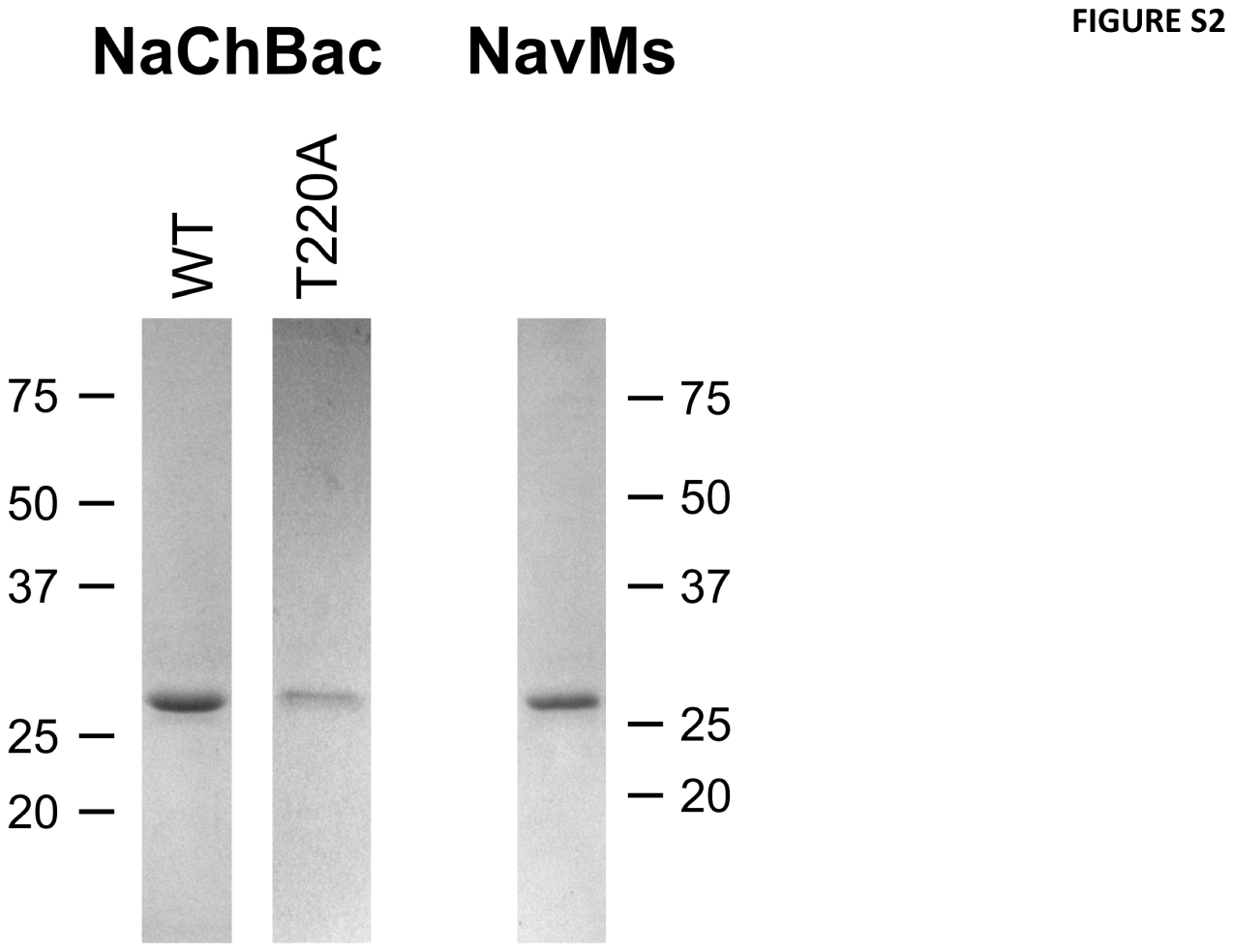

**Figure S2. Coomassie-stained SDS PAGE gels for purified NaChBac and NavMs proteins.** Positions of molecular weight standards are shown at either side.

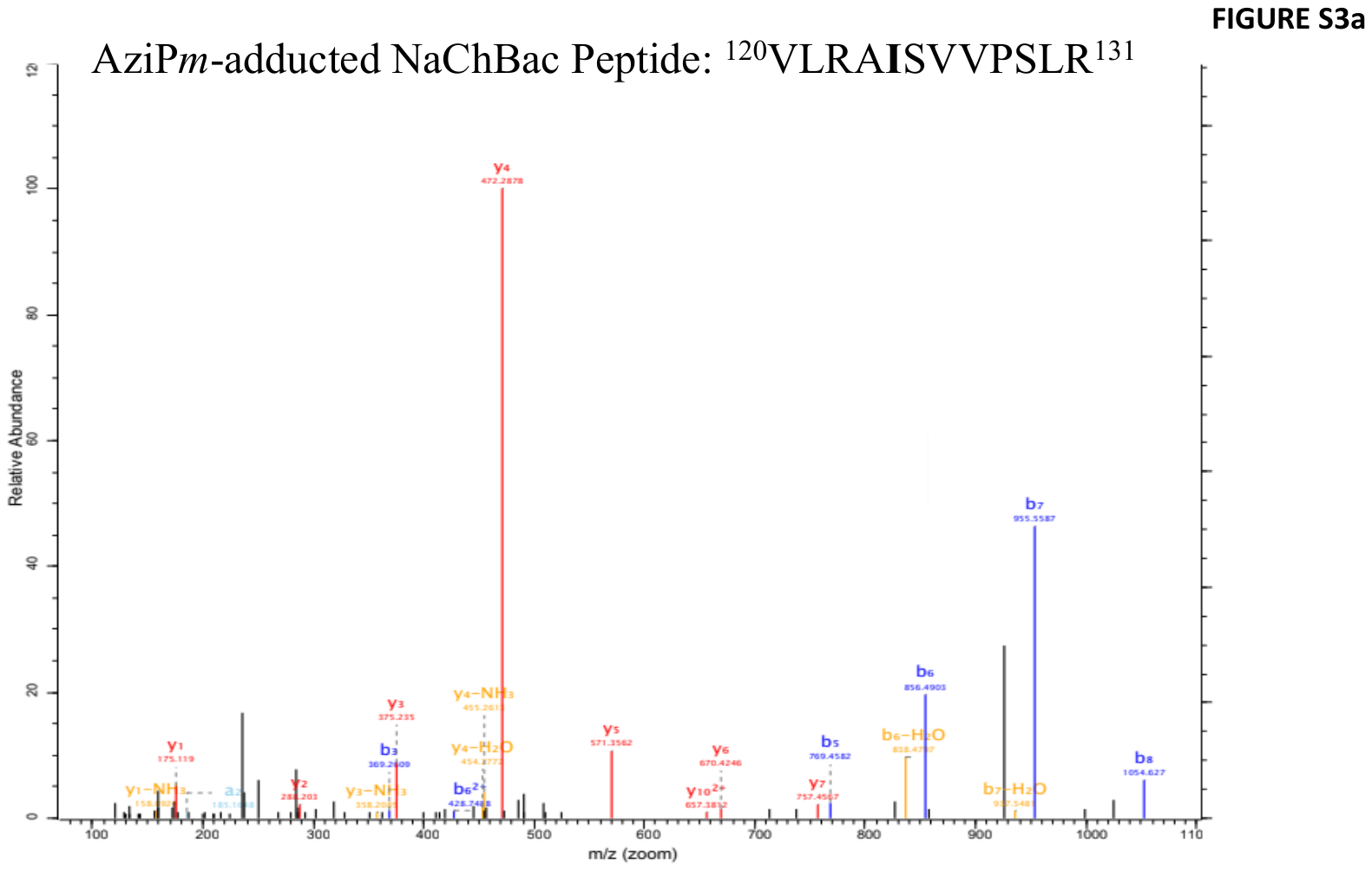

**Figure S3a. Mass spectra of AziP*m*-photolabeled peptides in NaChBac WT.** Identified a (light blue), b (blue), b-H_2_O(yellow), y (red), y-H_2_O (yellow), and Y-NH_3_(yellow) ions are labeled accordingly. Residues detected with an AziP*m* photomodification are bolded. The sequences coverage in mass spectrometry analysis was 77.8%.

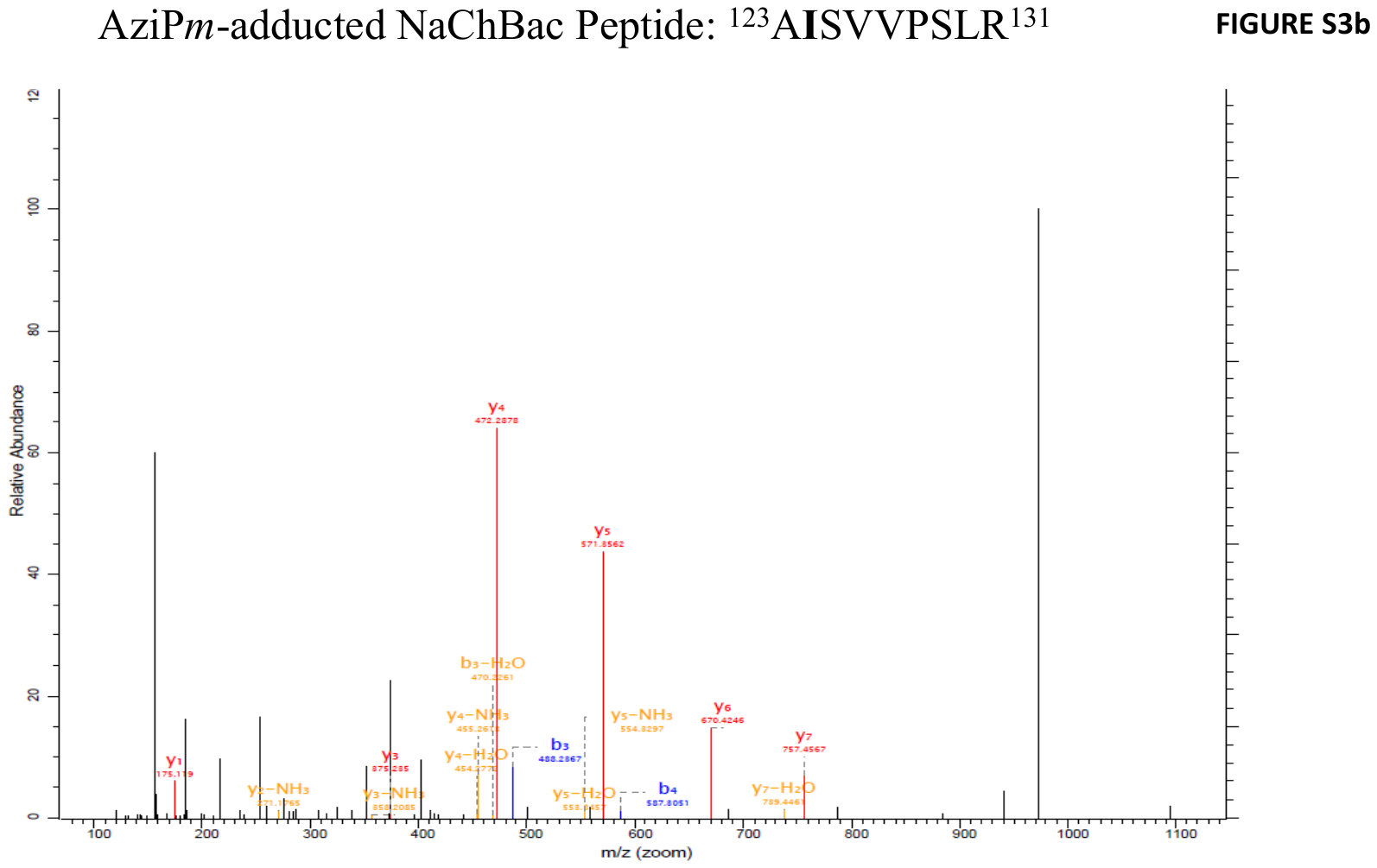

**Figure S3b. Mass spectra of AziP*m*-photolabeled peptides in NaChBac WT.** Identified a (light blue), b (blue), b-H_2_O(yellow), y (red), y-H_2_O (yellow), and Y-NH_3_(yellow) ions are labeled accordingly. Residues detected with an AziP*m* photomodification are bolded. The sequences coverage in mass spectrometry analysis was 81.6 %.

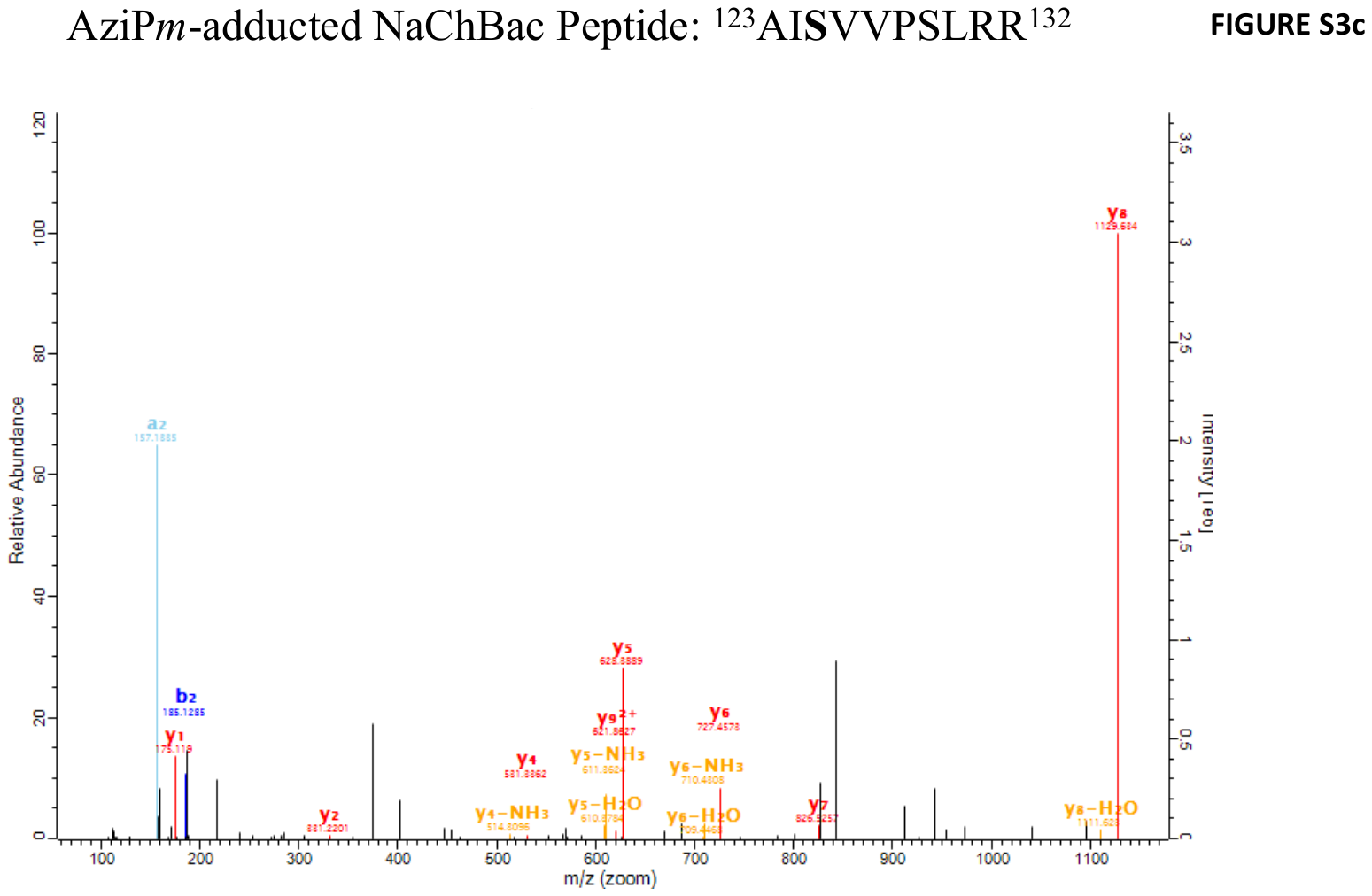

**Figure S3c. Mass spectra of AziP*m*-photolabeled peptides in NaChBac WT.** Identified a (light blue), b (blue), b-H_2_O(yellow), y (red), y-H_2_O (yellow), and Y-NH_3_(yellow) ions are labeled accordingly. Residues detected with an AziP*m* photomodification are bolded. The sequences coverage in mass spectrometry analysis was 79.6 %.

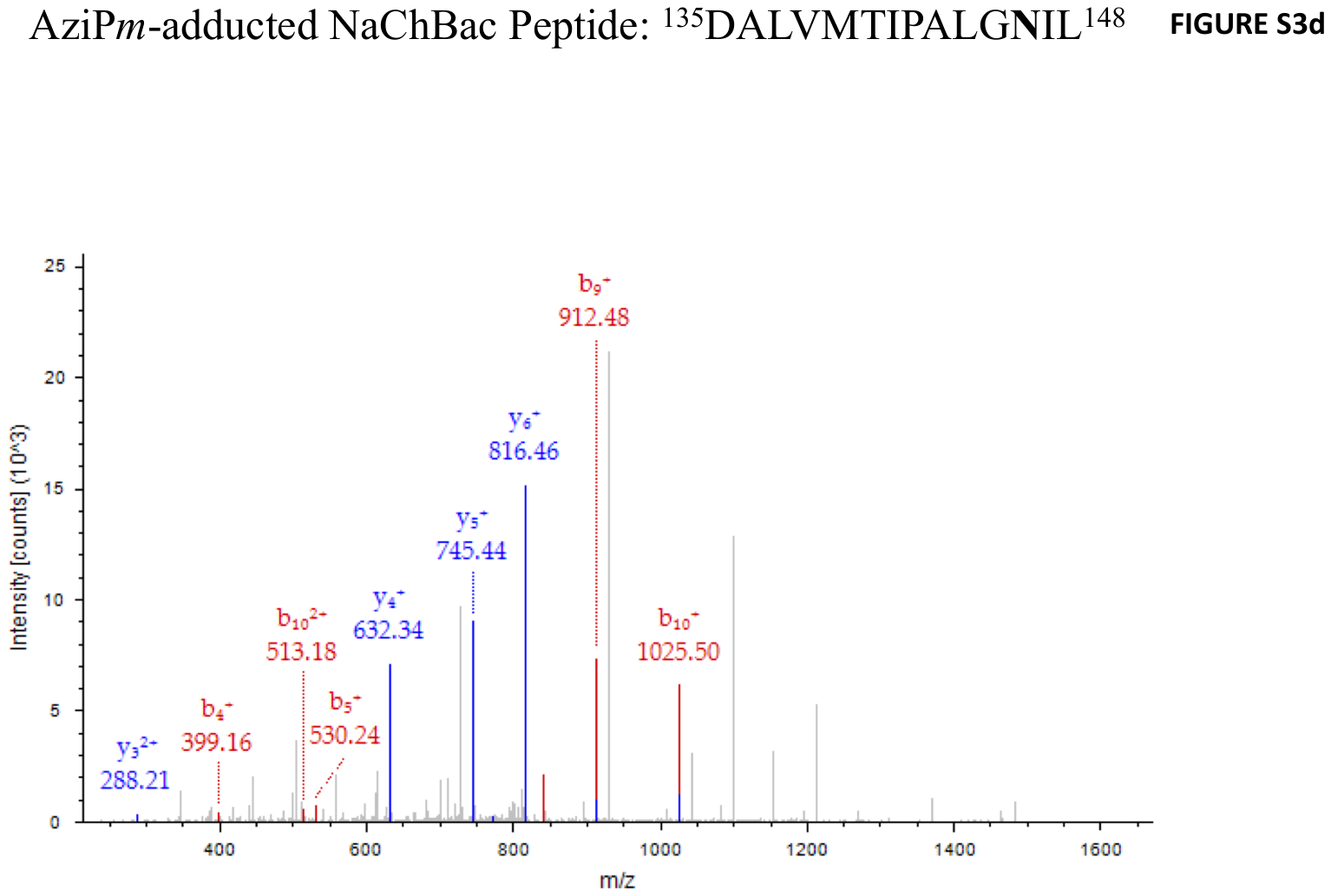

**Figure S3d. Mass spectra of AziP*m*-photolabeled peptides in NaChBac WT.** Identified b (red) and y (blue) ions are labeled accordingly. Residues detected with an AziP*m* photomodification are bolded. The sequences coverage in mass spectrometry analysis was 75.3 %.

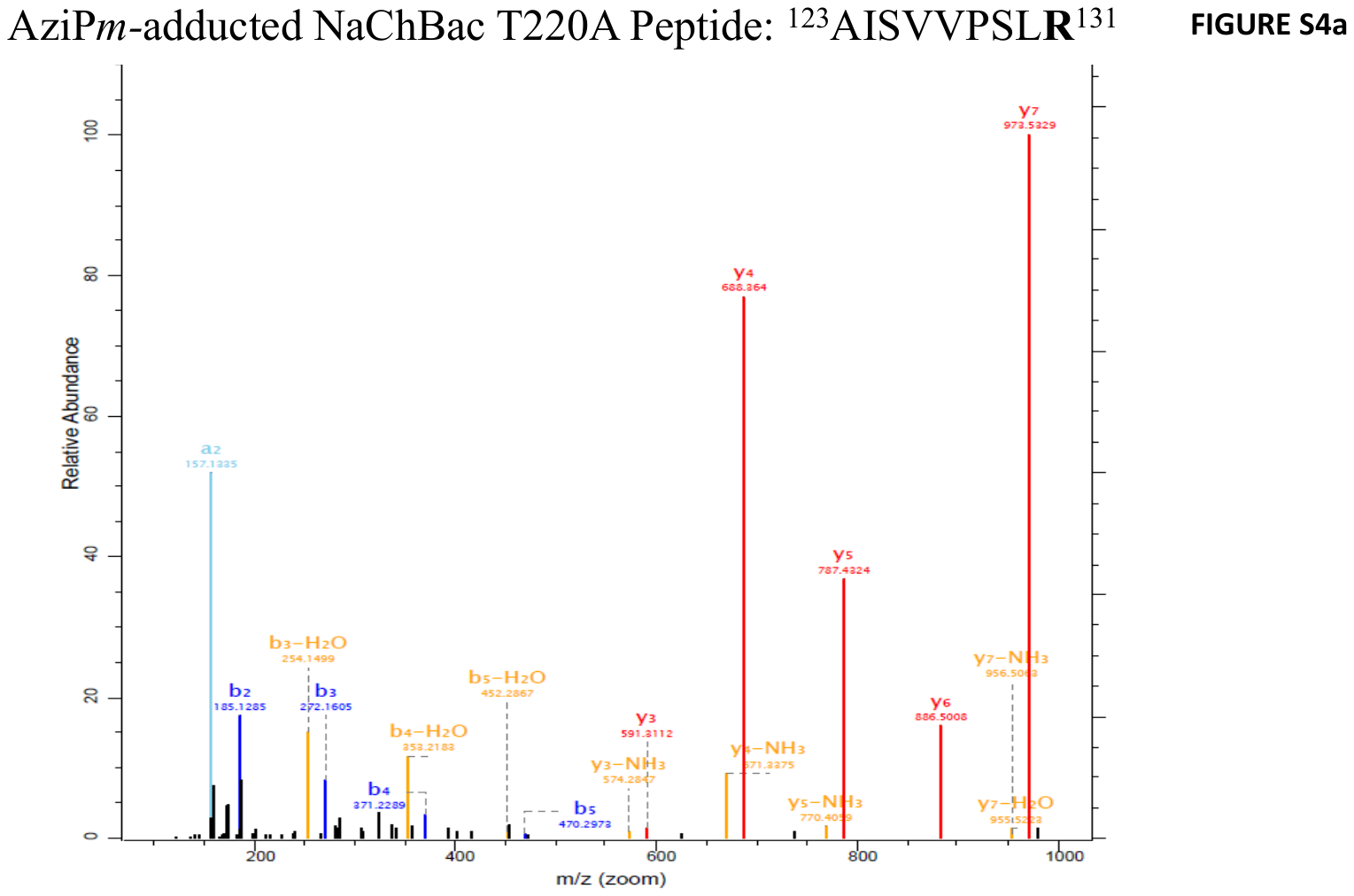

**Figure S4a. Mass spectra of photolabeled peptides in NaChBac T220A.** Identified a (light blue), b (blue), b-H_2_O(yellow), y (red), y-H_2_O (yellow), and Y-NH_3_(yellow) ions are labeled accordingly. Residues detected with an AziP*m* photomodification are bolded. The sequences coverage in mass spectrometry analysis was 75.3 %.

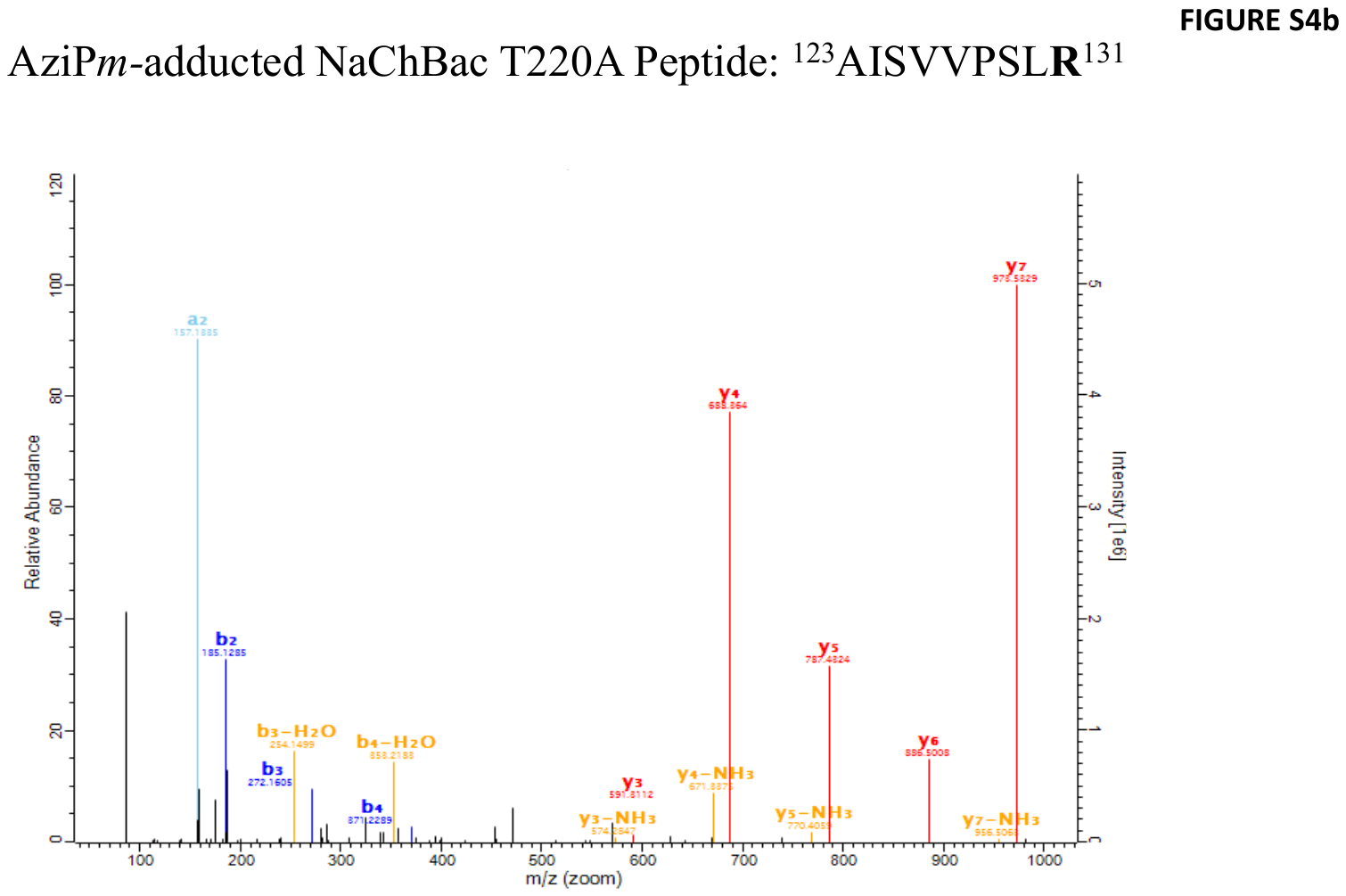

**Figure S4b. Mass spectra of photolabeled peptides in NaChBac T220A.** Identified a (light blue), b (blue), b-H_2_O(yellow), y (red), y-H_2_O (yellow), and Y-NH_3_(yellow) ions are labeled accordingly. Residues detected with an AziP*m* photomodification are bolded. The sequences coverage in mass spectrometry analysis was 76.9 %.

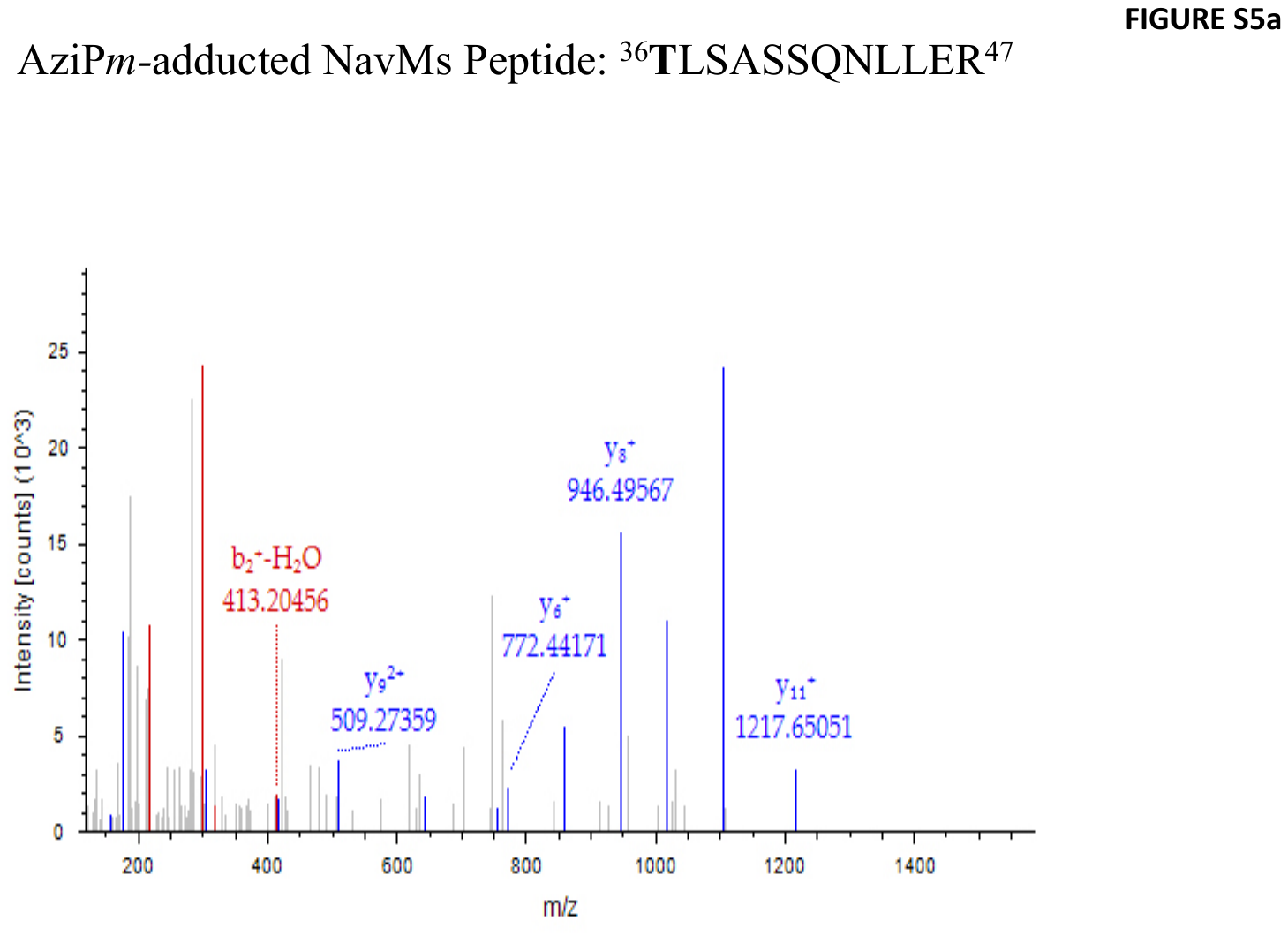

**Figure S5a. Mass spectra of photolabeled peptides in NavMs.** Identified b (red) and y (blue) ions are labeled accordingly. Residues detected with an AziP*m* photomodification are bolded. The sequences coverage in mass spectrometry analysis was 84.9 %.

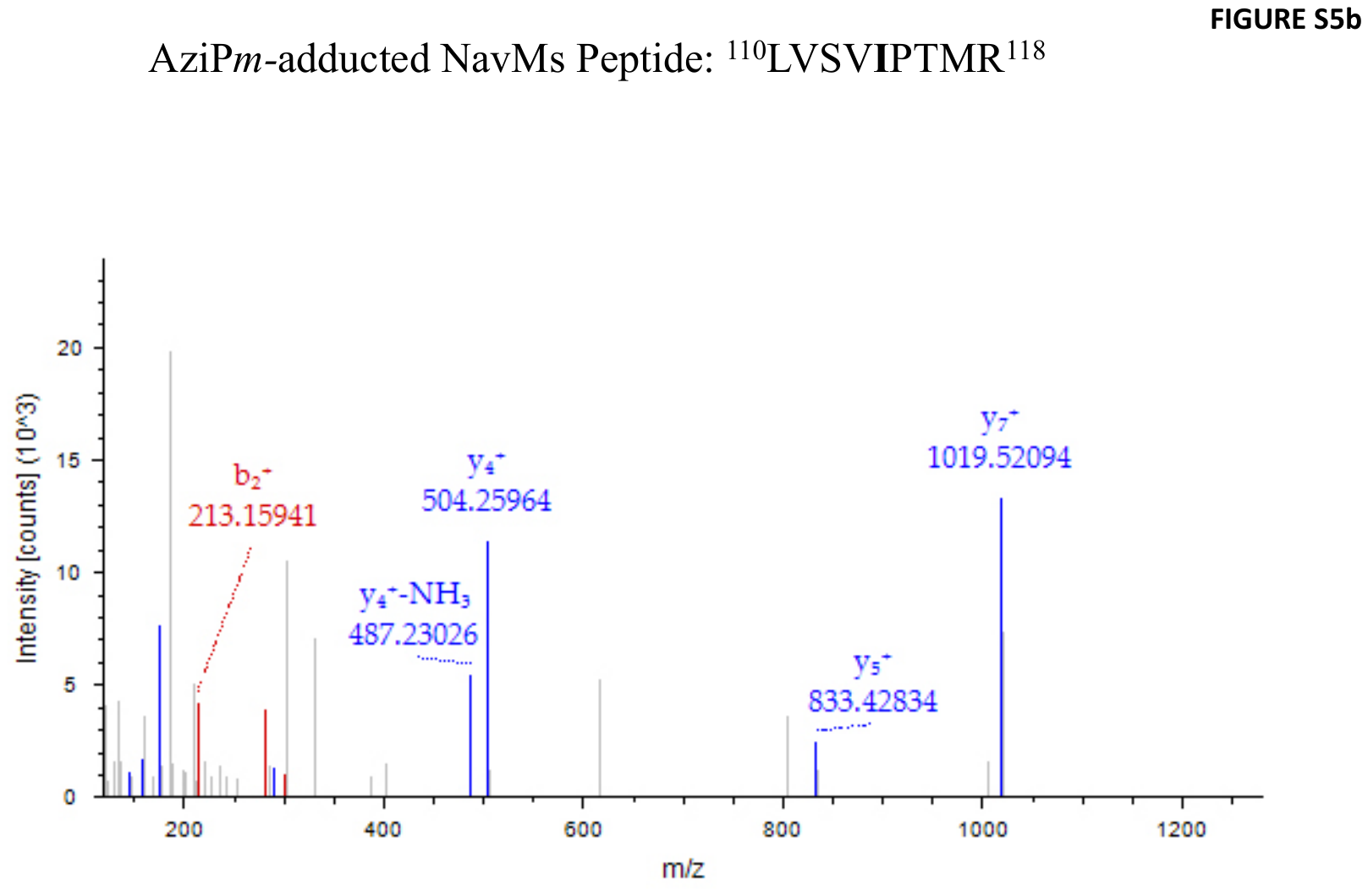

**Figure S5b . Mass spectra of photolabeled peptides in NavMs.** Identified b (red) and y (blue) ions are labeled accordingly. Residues detected with an AziP*m* photomodification are bolded. The sequences coverage in mass spectrometry analysis was 84.9 %.

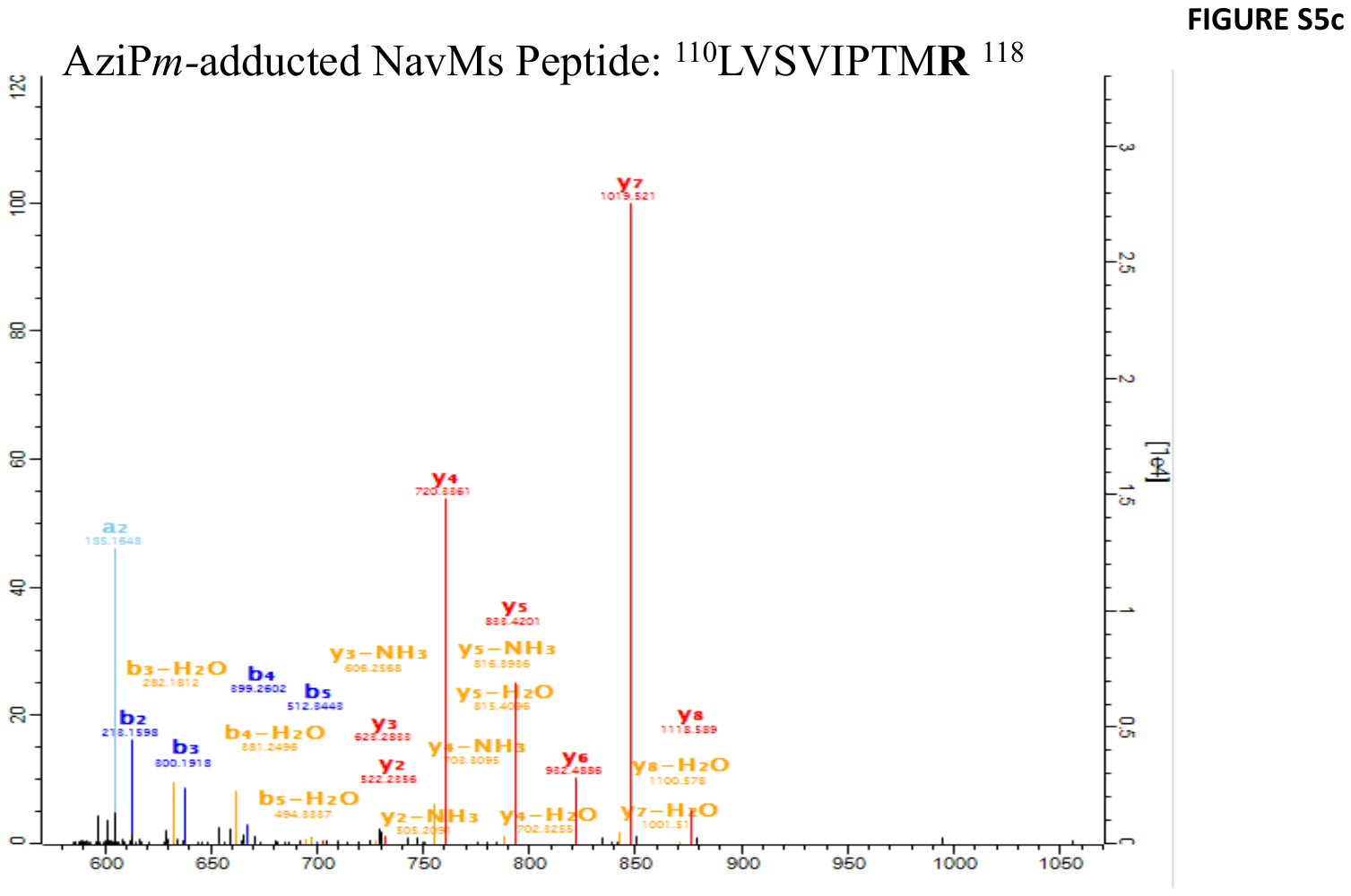

**Figure S5c. Mass spectra of photolabeled peptides in NavMs.** Identified a (light blue), b (blue), b-H_2_O(yellow), y (red), y-H_2_O (yellow), and Y-NH_3_(yellow) ions are labeled accordingly. Residues detected with an AziP*m* photomodification are bolded. The sequences coverage in mass spectrometry analysis was 45.8%.

His_6_-NaChBac WT

**MCHHHHHHGG SENLY**FQGM**K MEARQKQNSF TSKMQKIVNH RAF**TFTVIAL ILF**NALIVGI ETYPRIYADH KWLFYRIDLV LLWIFTIEIA MRFLASNPKS AFF**RSSWNWF **DFLIVAAGHI FAGAQFVTVL R**ILR**VLRVLR AISVVPSLRR LVDALVMTI**P ALG**N**ILILMS IFFY**IFAVIG TMLFQHVSPE YFGNLQLSLL TLFQVVTLES WASGVMRPIF AEVPWSWLYF VSFVLIG**TFI IFNLF**IGVIV NNVEKAELTD NEEDGEADGL KQEISALRKD VAELKSLLKQ SK**

His_6_-NaChBac-T220A

**MCHHHHHHGG SENLY**FQGMK MEARQK**QNSF TSKMQKIVNH R**AFTFTVIAL ILF**NALIVGI ETYPRIYADH KWLFYRIDLV LLWIFTIEIA MRFLASNPK**S AFFRSSWNWF DFLIVAAGHI F**AGAQFVTVL R**ILR**VLRVLR AISVVPSLRR LVDALVMTI**P ALGNILILMS IFFY**IFAVIG TMLFQHVSPE YFGNLQLSLL TLFQVVTLES WASGVMRPIF AEVPWSWLYF VSFVLIGA**FI IFNLF**IGVIV NNVEKAELTD NEEDGEADGL KQEISALRKD VAELKSLLKQ SK**

His_6_-NavMs

**MCHHHHHHGG SENLY**FQGMS RK**IRDLIESK RF**QNVITAII VLNGAVLGLL TDT**T**LSASSQ NLLER**VDQLC LTIFIVEISL KIYAYGVR**GF FRSGWNLFDF VIVAIALMPA QGSLSVLRTF **RIFRVMRLVS VIPTMRRVVQ GMLLALPGVG S**VAALLTVVF YIAAVMATNL YGATFPEWFG DLSKSLYTLF **QVMTLESWSM GIVR**PVMNVH PNAWVFFIPF IMLTTFTVLN LFIGIIVDAM AITK**EQEEEA KTGHHQEPIS QTLLHLGDRL DRIEKQLAQN NELLQR**QQPQ KK

**Figure S6. Sequence coverage map for NaChBac, NaChBac T220A, and NavMs mass spectrometry analysis.** Sequences with high confidence coverage in mass spectrometry analysis are bolded. AziP*m* adducted residues are noted in magenta, and the start methionine is indicated in green . Transmembrane domain helices are underlined, and the S4-S5 linker is highlighted in grey.

```
                                   ----------S1-----------       --------
NavMs        -----------MSRKIRDLIESKRFQNVITAIIVLNGAVLGLLTDTTLSASSQNLLERVD
NavAb        -----------MYLRITNIVESSFFTKFIIYLIVLNGITMGLETSKTFMQSFGVYTTLFN
NaChBac      MKMEARQKQNSFTSKMQKIVNHRAFTFTVIALILFNALIVGIETYPRIYADHKWLFYRID
                        :  :: .:::   *   :  :*::*.  :*: *   :  .       .:


             -----S2--------            -------S3------        ----------
NavMs        QLCLTIFIVEISLKIYAYG-VRGFFRSGWNLFDFVIVAIALMPAQG-SLSVLRTFRIFRV
NavAb        QIVITIFTIEIILRIYVHR--ISFFKDPWSLFDFFVVAISLVPTSS-GFEILRVLRVLRL
NaChBac      LVLLWIFTIEIAMRFLASNPKSAFFRSSWNWFDFLIVAAGHIFAGAQFVTVLRILRVLRV
              : : ** :** ::: .     .**:. *. ***.:** . : : .  . :** :*::*:

             --S4--- ---Linker---  ----------S5------------           ---
NavMs        MRLVSVIPTMRRVVQGMLLALPGVGSVAALLTVVFYIAAVMATNLYGATFPEWFGDLSKS
NavAb        FRLVTAVPQMRKIVSALISVIPGMLSVIALMTLFFYIFAIMATQLFGERFPEWFGTLGES
NaChBac      LRAISVVPSLRRLVDALVMTIPALGNILILMSIFFYIFAVIGTMLFQHVSPEYFGNLQLS
             :* ::.:* :*::*..:: .:*.: .:  *:::.*** *::.* *:    **:** *  *

             ---P1--- SF  ----P2----- -----------S6----------------
NavMs        LYTLFQVMTLESWSMGIVRPVMNVHPNAWVFFIPFIMLTTFTVLNLFIGIIVDAMAITKE
NavAb        FYTLFQVMTLESWSMGIVRPLMEVYPYAWVFFIPFIFVVTFVMINLVVAIIVDAMAILNQ
NaChBac      LLTLFQVVTLESWASGVMRPIFAEVPWSWLYFVSFVLIGTFIIFNLFIGVIVNNVEKAEL
             : *****:*****: *::**::   * :*::*: *::: ** ::**.:.:**: :   :


NavMs        QEEEAKTG---HHQEPISQTLLHLGDRLDRIEKQLAQNNELLQRQQPQKK
NavAb        KEEQHIIDEVQSHEDNINNEIIKLREEIVELK-------ELIKTSLKN--
NaChBac      TDN-----EEDGEADGLKQEISALRKDVAELKSLLKQSK-----------
              ::         . : :.: :  * . : .::
```

**Figure S7:**  **Sequence Alignments of NaChBac, NavAb and NavMs**. Adducted residues in NaChBac and NavMs are bolded and indicated in magenta. Transmembrane domain (S1-S6), S4-S5 linker, Pore region, the selectivity filter (SF) are indicated.

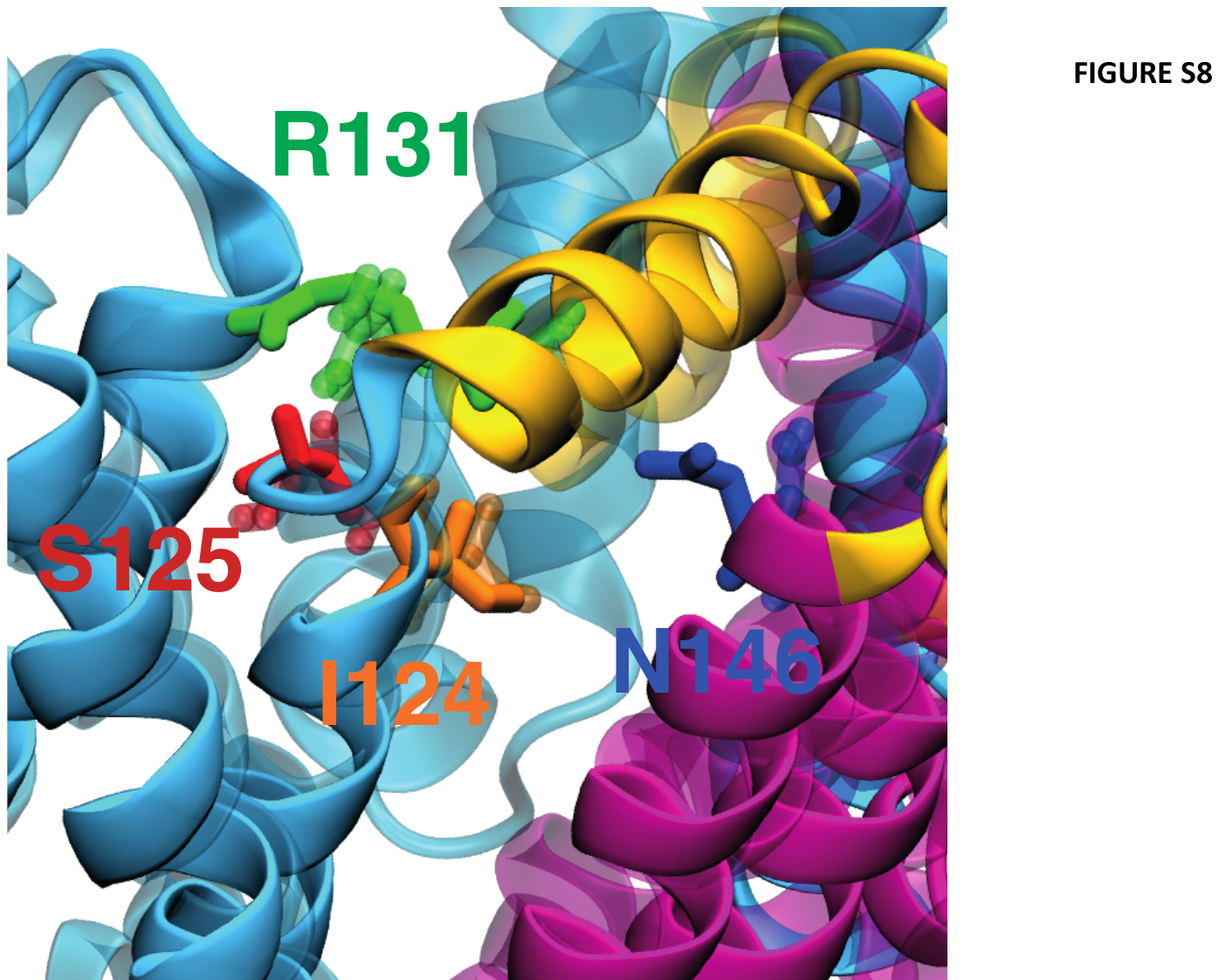

**Figure S8. Overlay of model and cryoEM structures of NaChBac.** The structure of of the theoretical model used in this study and the experimental cryoEM structure of NaChBac are shown superimposed. Both structures are shown as ribbons, with the former rendered in solid colors and the latter as shaded transparency. The sidechains of residues I124, S125, R131, and N146 are shown as sticks for both structures. Note that the two models are coincident and that the rotameric state of the residues analyzed in this paper is the same.

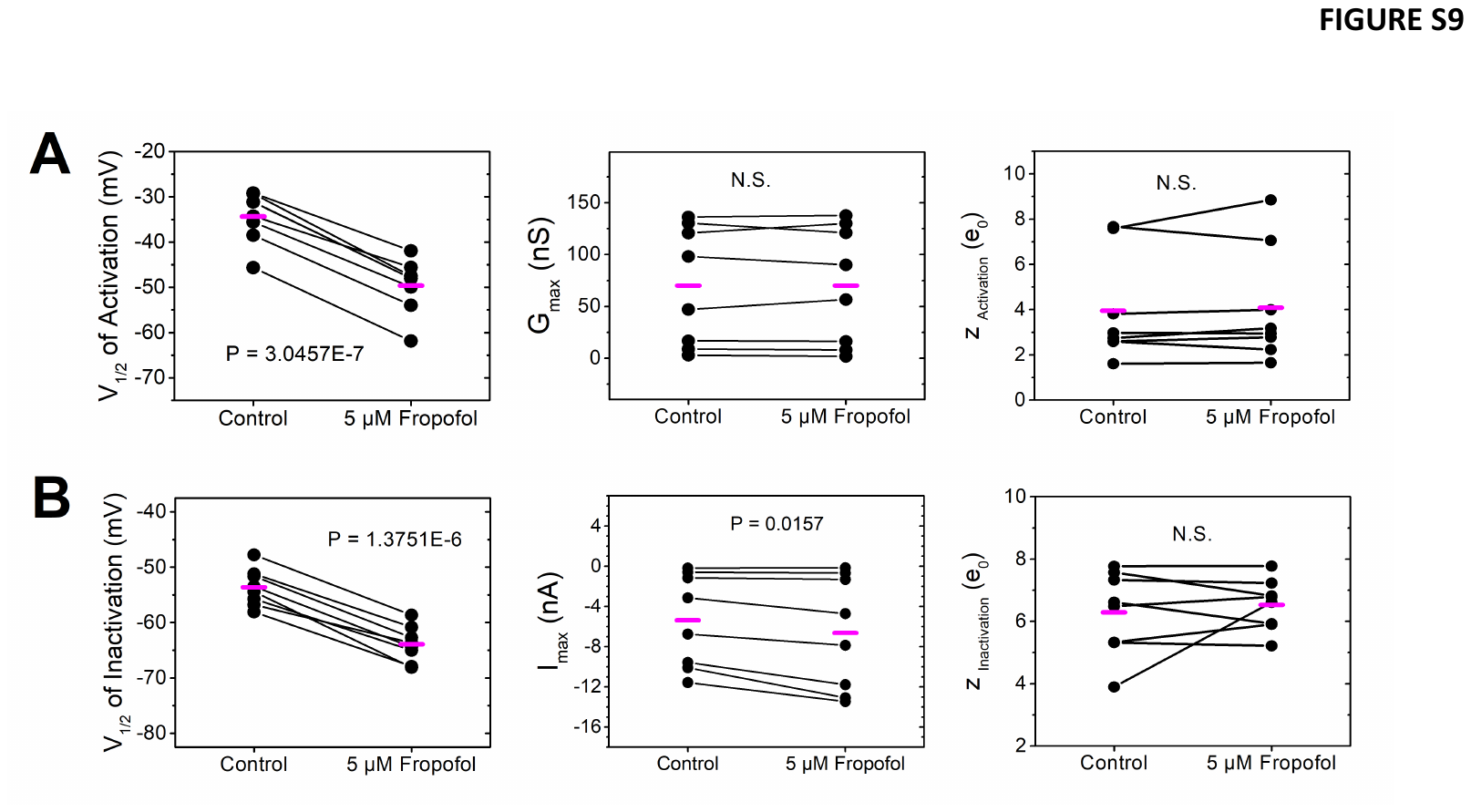

**Figure S9. Statistical analysis of changes in NachBac gating parameters induced by 5 μM fropofol**. A) Scatter graphs of V_1/2_, G_max_ and z derived from the best-fit to 4^th^-order Boltzmann functions (Materials and Methods). B) Scatter graphs of V_1/2_, I_max_ and z derived from the best-fit to 1^st^ order Boltzmann functions (Materials and Methods). The paired Student’s t-test was used to evaluate the changes. The p values are indicated in each graph (N.S. = not significant). **Source Data 6-8**

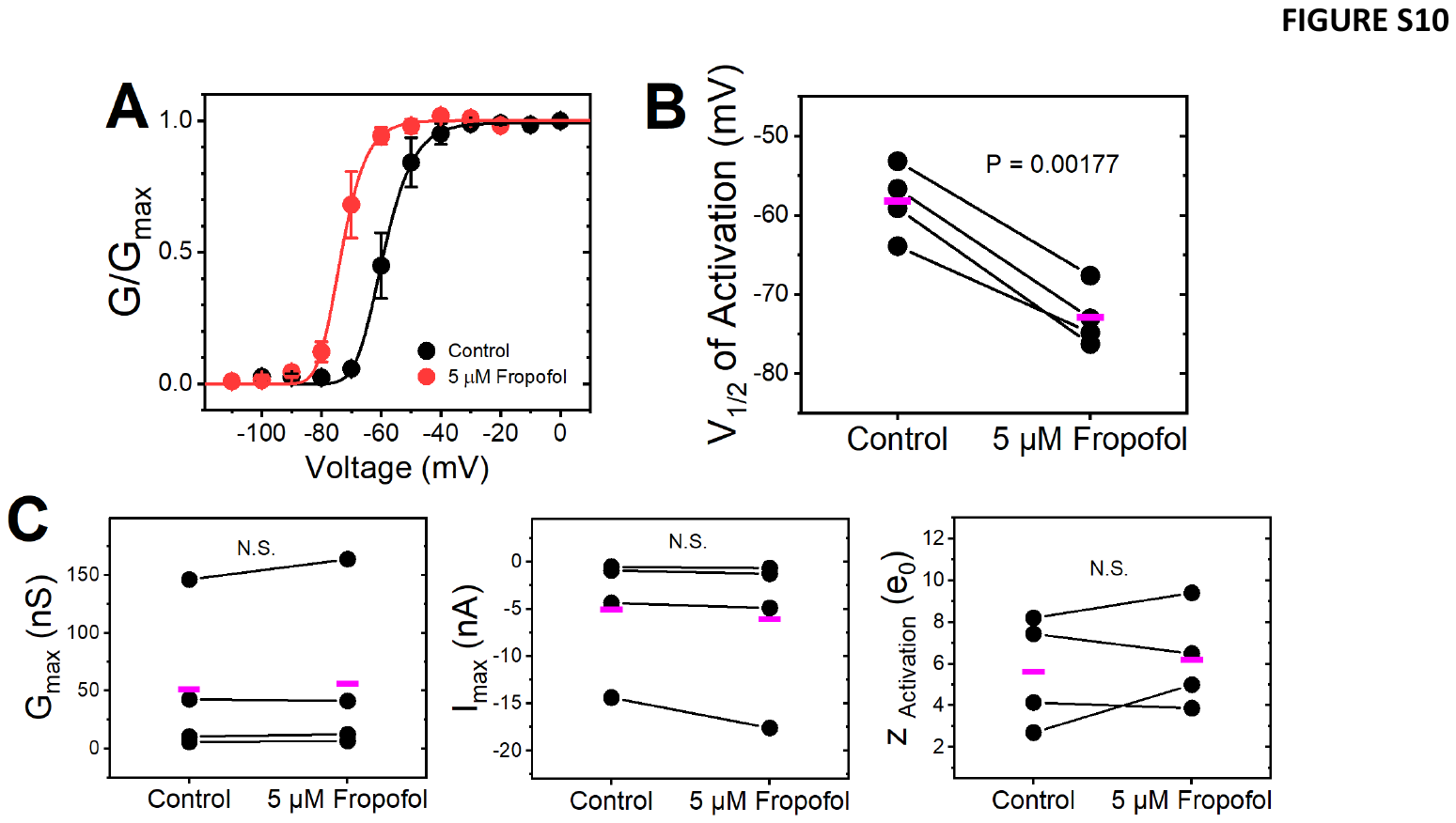

**Figure S10. Modulation of NaChBac T220A by fropofol.** A) Normalized peak conductance-voltage relations. The solid lines are best-fit 4^th^ order Boltzmann functions (summary of best-fit parameters on Table 1). B) Statistical analysis of changes in gating parameters induced by 5 μM fropofol. The paired Student’s t-test was used to evaluate the changes. The p values are indicated in each graph (N.S. = not significant). **Source Data 9-10.**

**SOURCE DATA (TO BE PROVIDED UPON REQUEST)**

Source Data 1: (WT + AziPm) values used to generate G/Gmax and I/Imax curves, tau inactivation; values for V1/2 (activation and inactivation), Gmax, z (activation and inactivation), Imax

Source Data 2: (WT + Fropofol) values used to generate G/Gmax and I/Imax curves, tau inactivation; values for V1/2 (activation and inactivation), Gmax, z (activation and inactivation), Imax

Source Data 3: (T220A + Fropofol) values used to generate G/Gmax curves; values for V1/2 activation, Gmax, z activation

Source Data 4: Values used to generate normalized I-V curves (WT + AziPm, WT + Fropofol, T220A + Fropofol)

Source Data 01: (WT + Control) activation currents

Source Data 02: (WT + AziPm 1 uM) activation currents

Source Data 03: (WT + Control) activation currents

Source Data 04: (WT + AziPm 5 uM) activation currents

Source Data 05: (WT + Control) inactivation currents

Source Data 06: (WT + AziPm 1 uM) inactivation currents

Source Data 07: (WT + Control) inactivation currents

Source Data 08: (WT + AziPm 5 uM) inactivation currents

Source Data 09: (WT + AziPm) values used to generate G/Gmax and I/Imax curves, tau inactivation; values for V1/2 (activation and inactivation), Gmax, z (activation and inactivation), Imax

Source Data 10: (WT + Control) activation currents

Source Data 11: (WT + Fropofol) activation currents

Source Data 12: (WT + Control) inactivation currents

Source Data 13: (WT + Fropofol) inactivation currents

Source Data 14: (WT + Fropofol) values used to generate G/Gmax and I/Imax curves, tau inactivation; values for V1/2 (activation and inactivation), Gmax, z (activation and inactivation), Imax

Source Data 15: (T220A + Control) activation currents

Source Data 16: (T220A + Fropofol) activation currents

Source Data 17: (T220A + Fropofol) values used to generate G/Gmax curves; values for V1/2 activation, Gmax, z activation, rising phase tau

Source Data 18: MD stuff?

Source Data 19: Values used to generate normalized I-V curves (WT + AziPm, WT + Fropofol, T220A + Fropofol)

**DATA LABELS IN SOURCE DATA**

Source Data 01/02: NaChBac WT Activation 1 uM AziPm

- 9/12/2017 Cell 6	17912001/17912003	1
- 9/12/2017 Cell 7	17912004/17912006	2
- 9/19/2017 Cell 2	17919000/17919004	3
- 9/19/2017 Cell 3	17919005/17919007	4
- 9/19/2017 Cell 4	17919009/17919012	5
- 9/21/2017 Cell 2	17921000/17921002	6
- 9/21/2017 Cell 6	17921004/17921006	7

Source Data 03/04: NaChBac WT Activation 5 uM AziPm

- 9/29/2017 Cell 1	17929000/17929002	8
- 9/29/2017 Cell 2	17929004/17929006	9
- 10/6/2017 Cell 3	17o06001/17o06003	10
- 10/6/2017 Cell 4	17o06005/17o06007	11
- 10/6/2017 Cell 11	17o06012/17o06013	12
- 10/10/2017 Cell 3	17o10000/17o10002	13
- 10/10/2017 Cell 4	17o10004/17o10006	14

Source Data 05/06: NaChBac WT Inactivation 1 uM AziPm

- 9/12/2017 Cell 6	17912002/17912004	15
- 9/14/2017 Cell 2	17914001/17914002	16
- 9/19/2017 Cell 2	17919001/17919003	17
- 9/19/2017 Cell 3	17919006/17919008	18
- 9/19/2017 Cell 4	17919011/17919013	19
- 9/21/2017 Cell 2	17921001/17921003	20
- 9/21/2017 Cell 6	17921005/17921007	21

Source Data 07/08: NaChBac WT Inactivation 5 uM AziPm

- 9/29/2017 Cell 1	17929001/17929003	22
- 9/29/2017 Cell 2	17929005/17929007	23
- 10/6/2017 Cell 3	17o06002/17o06004	24
- 10/6/2017 Cell 11	17o06011/17o06015	25
- 10/10/2017 Cell 3	17o10001/17o10003	26
- 10/10/2017 Cell 4	17o10005/17o10007	27

Source Data 10/11 NaChBac WT Activation Fropofol

- 10/26/16 Cell 4	16o26003/16o26005	28
- 10/27/16 Cell 5	16o27007/16o27009	29
- 12/13/16 Cell 1	16d13000/16d13002	30
- 12/13/16 Cell 2	16d13005/16d13007	31
- 12/13/16 Cell 3	16d13008/16d13010	32
- 12/14/16 Cell 2	16d14005/16d14007	33
- 12/14/16 Cell 4	16d14010/16d14012	34
- 12/14/16 Cell 5	16d14013/16d14015	35

Source Data 12/13: NaChBac WT Inactivation Fropofol

- 10/26/16 Cell 4	16o26004/16o26006	36
- 10/27/16 Cell 5	16o27008/16o27010	37
- 12/13/16 Cell 1	16d13001/16d13003	38
- 12/13/16 Cell 2	16d13006/16d13008	39
- 12/13/16 Cell 3	16d13009/16d13011	40
- 12/14/16 Cell 2	16d14006/16d14008	41
- 12/14/16 Cell 4	16d14011/16d14013	42
- 12/14/16 Cell 5	16d14014/16d14016	43

Source Data 15/16: NaChBac T220A Activation Fropofol

- 1/16/17 Cell 3	17116003/17116004	44
- 1/16/17 Cell 4	17116005/17116007	45
- 1/16/17 Cell 5	17116007/17116008	46
- 1/17/17 Cell 4	17117005/17117006	47
